# CoDIAC: A comprehensive approach for interaction analysis reveals novel insights into SH2 domain function and regulation

**DOI:** 10.1101/2024.07.18.604100

**Authors:** Alekhya Kandoor, Gabrielle Martinez, Julianna M. Hitchcock, Savannah Angel, Logan Campbell, Saqib Rizvi, Kristen M. Naegle

## Abstract

Protein domains are conserved structural and functional units that serve as building blocks of proteins. Through evolutionary expansion, domain families are represented by multiple members in diverse configurations with other domains, evolving new specificities for their interacting partners. Here, we develop a structure-based interface analysis to comprehensively map domain interfaces from experimental and predicted structures, including interfaces with macromolecules and intraprotein interfaces. We hypothesized that comprehensive contact mapping of domains could yield new insights into domain selectivity, conservation of domain-domain interfaces across proteins, and identify conserved post-translational modifications (PTMs), relative to interaction interfaces, allowing for the inference of specific effects due to PTMs or mutations. We applied this approach to the human SH2 domain family, a modular unit central to phosphotyrosine-mediated signaling, identifying a novel approach to understanding binding selectivity and evidence of coordinated regulation of SH2 domain binding interfaces by tyrosine and serine/threonine phosphorylation and acetylation. These findings suggest multiple signaling systems can regulate protein activity and SH2 domain interactions in a coordinated manner. We provide the extensive features of the human SH2 domain family and this modular approach as an open source Python package for COmprehensive Domain Interface Analysis of Contacts (CoDIAC).

## Introduction

Protein domain families share a common fold and function and serve as key building blocks for proteins. Recently, it was predicted that the human proteome contains 47,576 domains (1) with diverse functions including binding to proteins, RNA, DNA, or lipids, and enzymatic activities such as regulating phosphorylation, proteolysis, and DNA repair. Recombination of domains into new protein architectures can tailor existing functions or create entirely new ones. As domains duplicate and evolve, they develop new specificities, enabling novel functionality in both the domain-containing protein and its interaction partners (2). Despite learning much from individual domain components, we still lack complete understanding of how domain interaction interfaces drive molecular interactions and regulate overall protein function.

Domains are targets of numerous post-translational modifications (PTMs) (3), most with unknown functions. Previously, we hypothesized that conserved PTMs—those occurring in the same structural position across many domains in a family—likely share common effects. Applying this concept to the RRM domain family, we identified conserved tyrosine phosphorylation in the RNA binding motif, suggesting global regulation of RNA interactions by phosphotyrosine signaling (4). However, this approach lacked comprehensive analysis between PTMs and the myriad interaction interfaces across domains. Understanding residue-level contacts with ligands or between domains in a protein could help hypothesize functional effects of PTMs. While the Protein Contact Atlas (5) provided non-covalent interactions from X-ray structures, challenges connecting structures with reference sequences, lack of programmatic access, and absence of coverage for NMR, EM, and AlphaFold structures (6) led us to develop a modular Python-based package to extract interaction interfaces of domains from a wider variety of sources. This tool identifies relevant structures, defines domain boundaries, and integrates contact analysis with PTMs and mutations using flat text formats and Jalview-based visualization. Most modules can be used individually or together for comprehensive domain family analysis.

Given domains’ central role in protein function, this pipeline should enable diverse research applications. We apply it here to analyze SH2 domains—”reader” domains essential to phosphotyrosine (pTyr) signaling. The SH2 domain is interesting, because it is extensive (119 domains in the human proteome on 109 proteins), it co-occurs in diverse protein architectures (e.g alone, with other reader domains, and with a broad range of enzymatic domains), it is a member of a broad class of globular domains that interact with flexible protein ligands (termed domain-motif interactions), and it is extensively post-translationally modified. Despite considerable experimental coverage in the PDB, structures represent only a fraction of the “SH2ome”, with even fewer structures complexed with ligands. While progress has been made in understanding binding determinants, including developing SH2 domains as reagents for pTyr site enrichment in mass spectrometry (7), we still lack comprehensive understanding of which SH2 domains interact with which of the 46,000 pTyr sites in the human proteome (8). Better understanding could improve SH2 domain design as affinity reagents, while engineering new specificities for inhibitors targeting dysregulated tyrosine kinase signaling (9, 10). Additionally, though one structural position of tyrosine phosphorylation has a known regulatory function in SRC family kinases (11–13), little is known about other extensive PTMs on SH2 domains.

Here, we describe our generalized contact mapping and domain-centric analysis pipeline, COmprehensive Domain Interface Analysis of Contacts (CoDIAC), and its application to human SH2 domains. We developed an approach to systematically project contacts, including from AlphaFold predictions, to infer contact maps for SH2 domains lacking structural data. Our analysis revealed both known and novel insights. Through contact mapping, we found that the fraction of bonds formed between an SH2 domain and specific ligand positions directly correlates with the specificity conferred at those positions. We also discovered that domain-domain pairings recurring in the proteome maintain conserved interaction interfaces, but primarily when the entire protein architecture is conserved. Additionally, we found that partial protein structures sometimes misrepresent native interaction interfaces. Through conserved structural analysis of PTMs and binding interfaces, we identified extensive modification of both domain-domain and domain-ligand interfaces by serine/threonine phosphorylation, tyrosine phosphorylation, and acetylation, suggesting regulation of SH2 domain “reader” function by multiple signaling systems. Finally, our ability to extrapolate across the SH2 domain family while considering multifaceted interfaces provides new insights into clinically-relevant mutations.

## Results

### CoDIAC: A comprehensive domain-centric pipeline for analysis of contacts

CoDIAC is a Python-based package integrating multiple resources to identify and extract contact maps from all available experimental structures in the PDB (14) and AlphaFold predictions (6) that cover a domain of interest as defined by its InterPro accession (15). Figure 1A shows the CoDIAC pipeline, which builds an annotated reference set of proteins and structures, extracts contact maps, and integrates with other features to produce resource files for analysis and Jalview visualization (16). CoDIAC uses UniProt (17) as the primary protein reference and pairwise sequence alignment from Biopython (18) to align structural sequences with UniProt reference sequences, identifying domain boundaries and creating features referenced to a common sequence. Beyond structurally extracted contact maps, CoDIAC translates other protein annotations onto reference sequences and integrates extraction of PTMs from ProteomeScout (3) and PhosphoSitePlus (19), as well as variants from databases like OMIM (20) and clinically significant gnomAD missense mutations (21, 22). CoDIAC is available at https://github.com/NaegleLab/CoDIAC.

**Fig. 1.**
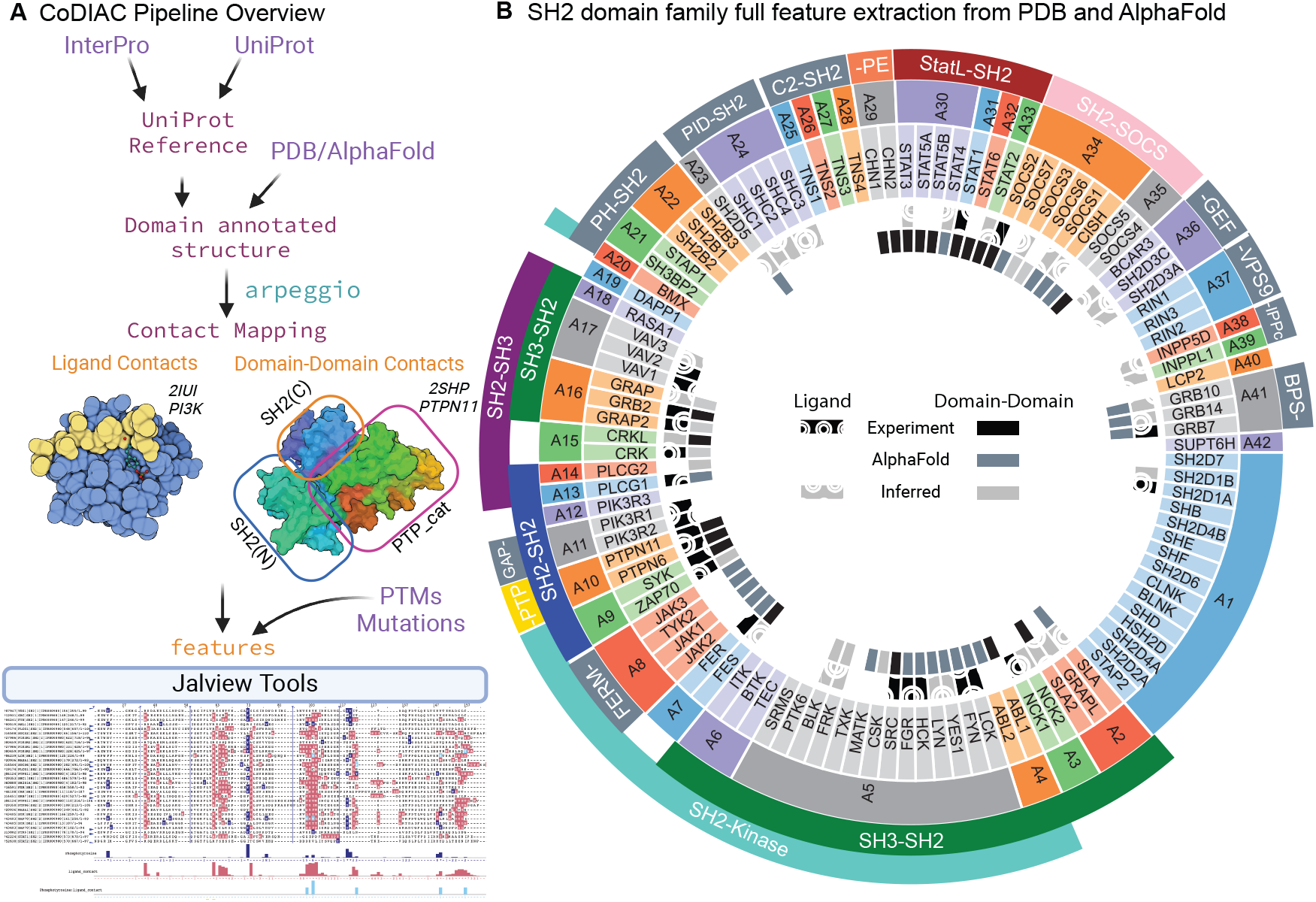
**A)** Flowchart outlining the key modules of the CoDIAC pipeline that can identify relevant domain containing proteins (agnostic of species), associated PDB and AlphaFold structures, annotate controlled domains from those resources for the extraction of domain interactions between proteins (ligand) and within proteins (domaindomain), and orient PTMs, mutations, and contact interfaces onto a common reference and Jalview-based visualization. **B)** An overview of all data and features extracted for the human SH2 domain family using CoDIAC. The SH2 domain family has been sorted and grouped by shared architectures (A1 for example is a group that contain only the SH2 domain). The immediately adjacent domains to the SH2 domain are labeled on the outermost tracks (e.g. A5 group is an architecture that consists of SH3-SH2-Kinase) and are in color if experimental structures covering the pair of domains exists. The feature generation and inference performed for the SH2 domain family are highlighted in the inner two circles. The innermost circle represents the genes for which we have experimental (black), AlphaFold (slate-gray), and inferred (light gray) features for domain-domain interfaces and the outer circle with hatched pattern represents ligand interface contacts with phosphotyrosine (pTyr)-containing ligands.

While we demonstrate domain-centric analysis of the human SH2 domain family using the complete CoDIAC pipeline, its modular components can be used independently. The first part produces descriptive text files containing se-quences, gene names, and domain annotations for proteins of interest, useful for evolutionary studies across species or comprehensive proteome information. The feature integration components can be used beyond contact mapping to merge different resources. For example, here we integrate manually extracted features from prior research on phage display mutagenesis (23) and SH2 domain phospholipid binding contacts (24). Additionally, the contact map extraction component is domain-agnostic and can analyze any regions within structures. This modular framework provides a flexible toolkit for various protein and domain-based analyses.

### CoDIAC contact mapping

A key aspect of CoDIAC is the conversion of structural files into fingerprints of non-covalent interactions. This enables domain-focused analysis by mapping contacts across all identified PDB and AlphaFold structures containing the domain of interest. For individual structures, CoDIAC uses Arpeggio (25) to generate flat text files of all interatomic interactions. These “adjacency files” contain distance and contact type information for each interacting residue pair across entities and chains. We then create binary adjacency files indicating whether pairwise residue-level interactions have sufficient evidence (binary value 1) or not (binary value 0), based on user-defined parameters for maximum interaction distance or contact types. For experimental structures with multiple molecule representations, we aggregate across assemblies to determine sufficient residue-level contact evidence (also user-defined). For SH2 domain analysis, we retained interactions with distances under 5Å and present in at least 25% of chains. For structures with multiple entities (domain and ligand), we kept contacts shared by at least 50% of domain-ligand pairs. The result is a contact map represented as a binary adjacency file detailing interactions with PDB ID, residue numbers, IDs, entity IDs, and binary interaction values. CoDIAC processes mmCIF files from any source, including PDB and AlphaFold, offering flexibility in contact mapping parameters within a common programming environment and producing readable file formats.

For domain-centric analysis, CoDIAC uses generated contact maps and annotated structure files to identify domain regions and analyze its interactions with other regions. In this work, we focused on SH2 domain interactions with other domains within a protein or with phosphotyrosine-containing ligands. CoDIAC creates Jalview-style feature files of contact maps for visualization and analysis. Since proteins often have multiple experimental structures, we aggregate across independent experiments to produce a single set of contact features for each available SH2 domain interaction interface. To avoid spurious contacts from study bias, we required features to be shared in at least 30% of available structures when multiple structures exist, a threshold determined by measuring feature retention versus inclusion stringency (Fig S1). The resulting pipeline produces feature files indicating which SH2 domain residues interact with ligands or other domains, enabling exploration of domain regulation by PTMs or mutations and their effects on protein function and ligand binding.

### SH2 domain analysis of interfaces

We used CoDIAC to systematically extract domain interfaces from structures to understand binding specificity, interface conservation, and the function of mutations and PTMs on SH2 domains. Figure 1B depicts the human SH2 domain family (119 domains across 109 proteins) with their protein architectures and interface coverage from experimental structures, AlphaFold predictions, or conservation-based inference. From the PDB, we identified 467 experimental structures covering 61 unique human SH2 proteins, with 135 structures including pTyrcontaining ligand complexes. Most ligand-containing structures (101) occur in trans (ligand and domain as separate entities), while 34 are bound in cis, with the ligand part of the same protein chain (Fig S2). For contact mapping, we excluded structures with variants in the SH2 domain during ligand interface mapping, and structures with any mutations during domain-domain interface mapping. The experimental structures provided 84 unique canonical SH2-pTyr ligand pairs and 207 structures across 32 proteins with domain-domain interface coverage, including 144 representing full protein architectures (Fig S3). Although AlphaFold currently cannot predict ligand-bound structures involving modified residues, we leverage the full length predicted structures of SH2 domain-containing proteins with low prediction error to study domain–domain interaction interfaces. We obtained 109 predicted structures for full protein architecture domain-domain interface analysis. We used PROMALS3D (26) to create a structure-based alignment of the SH2 domain family, with all data available at https://figshare.com/account/home#/projects/211453, representing a comprehensive resource of SH2 domain interfaces.

### Comprehensive ligand contact mapping

We extracted SH2 domain residues interacting with ligands across all available structures. Figure 2A shows residue-level contacts for 28 unique SH2 domains and their ligands from 102 PDB structures. Canonical SH2 binding involves a central engagement of the pTyr residue with an invariant arginine in the binding pocket (alignment position 62), with specificity from surrounding residues interacting within a shallow pocket (27). Approximately half the binding energy comes from the invariant arginine-pTyr interaction, with the remainder from nearby ligand residues (28). Our systematic analysis recovered the pTyr-invariant arginine interaction in most cases, with nine exceptions occurring under high local concentrations (Fig. S4), including tandem SH2 domains with bivalent pTyr ligands (SYK, ZAP70, RASA1) and cases where SH2 domains (CRK, CRKL) are tethered to pTyr ligands in cis. While these exceptions highlight possible non-canonical binding, we excluded them from canonical binding pocket mapping.

**Fig. 2.**
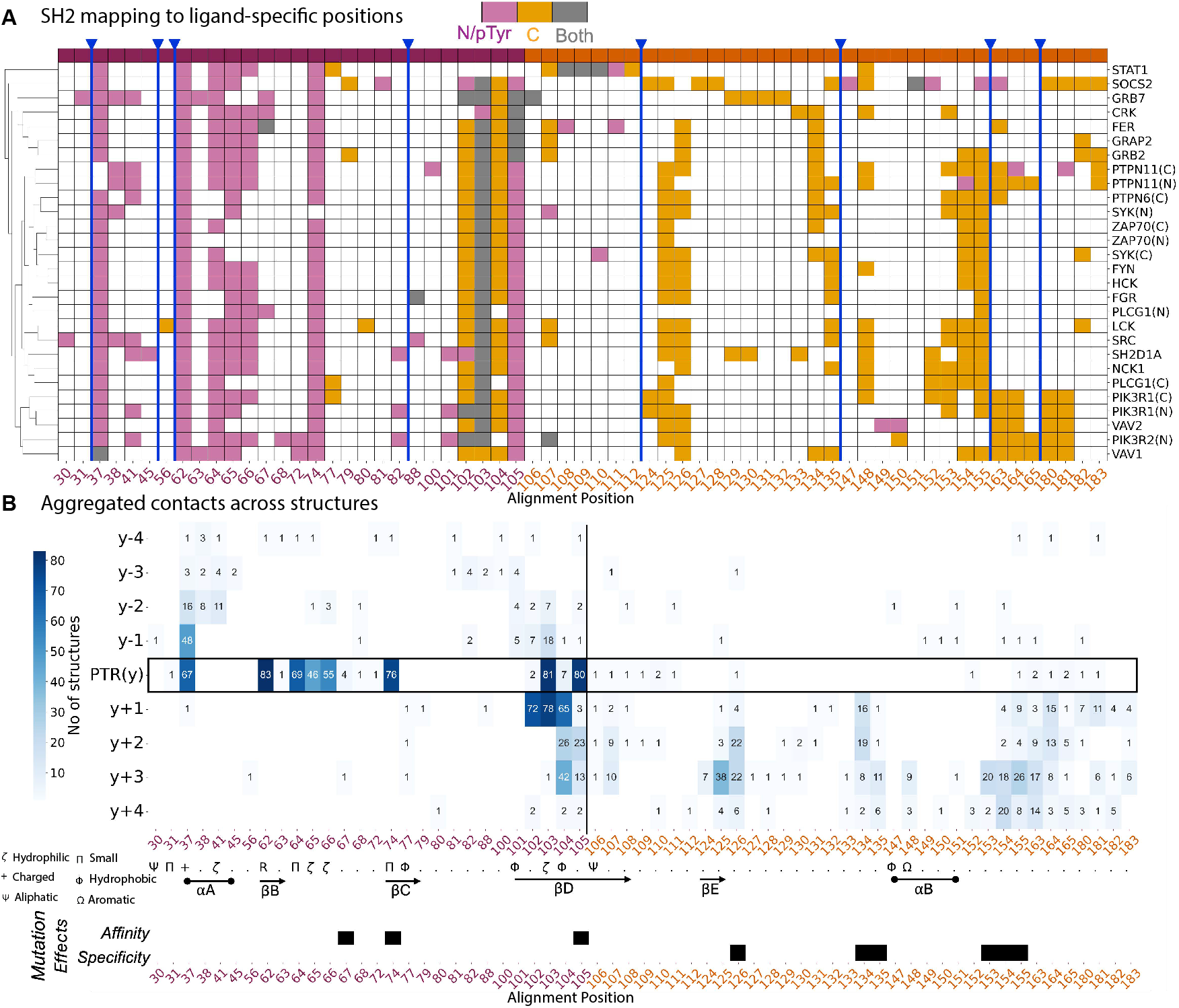
**A)** A matrix of ligand-based contacts of the SH2 domain relative to the reference alignment of the SH2 domain family (selected examples). Contacts are colored, based on if the residue makes contact to the N-terminal side or the pTyr on the ligand (pink), to the C-terminal side (orange), or to both sides (gray). Position 105 is the most C-terminal position we see consistently interact with the N-terminal portion of the ligand. **B)** Matrix representation of the combined feature set for canonical binding interface of SH2-pTyr interactions, relative to the reference alignment of SH2 domains and oriented by the pTyr for ligands. Features viewed here are aggregated from across 84 unique canonical SH2-pTyr ligand pairs and each feature is projected as an interaction between the residues on domain (x-axis) and ligand (y-axis). Secondary structures and amino acid consensus annotations are obtained from the PROMALS3D alignment. Bottom panel: Representative mutations and their effects from phage display studies separated according to whether they increase affinity (i.e. “superbinder”) or alter specificity.

We explored how important the sequence identity of the ligand was to the mapping of the SH2 domain ligand binding pocket. We encoded the SH2 domain residues that make contact with the ligands according to whether they make contact with the N-terminal or pTyr of the ligand, the C-terminal side of the ligand, or both N-term/pTyr and C-term (Fig. S5 and Fig. 2A). This recapitulated findings from individual structure studies, showing N-terminal SH2 regions engage N-terminal and pTyr ligand positions, while C-terminal SH2 binds C-terminal ligand positions. We identified position 105 in our reference alignment as the primary bifurcation point between pTyr pocket and specificity region across most domains (Fig. 2A). When we similarly encoded ligand residues by their interaction with SH2 domain regions before or after position 105, clustering showed ligand interactions are driven by SH2 domain identity rather than ligand sequence (Fig. S6), consistent with flexible ligands fitting into rigid globular domains.

We also investigated how ligand length affects domain mapping by examining residues mapped by just the pTyr (Fig. S7) and additional flanking residues (Fig. S5). We found the presence of N-terminal amino acids were unnecessary for recovering unique contacts, as ligands consistently mapped the same pocket residues regardless of N-terminal presence (see STAT1 and PIK3R1(C) domain binding in Fig. S5). Furthermore, even short ligands often mapped the same SH2 domain residues as longer ones, suggesting additional ligand residues frequently do not contribute novel domain contacts (see GRB2 in Fig. S5). However, in some cases, very short ligands only mapped part of the specificity region, while longer peptides revealed additional contacts (see SRC in Fig. S5). The high consistency of domain residues mapped across independent experiments with different ligand sequences and lengths suggests the available PDB ligands provide robust mapping of the SH2 domain binding pocket, especially when aggregating information across structures.

Next, we explored the interactions between specific SH2 domain residues and ligand positions. Figure 2B provides a 2D view with SH2 domain alignment positions on the x-axis, ligand positions on the y-axis, and values indicating the number of structures mapping contacts between position pairs. The most frequent interaction occurs between the invariant arginine (position 62) and the pTyr residue. High agreement also exists for other conserved structural positions interacting with the pTyr, including positions 103 (polar/His dominant) and 105 (hydrophobic/Lys dominant) on the *β*D-strand, which engage pTyr in >95% of structures, and position 37 on the *α*A helix (mostly arginine), which engages pTyr in 80% of structures. These enriched positions align with known SH2-pTyr binding mechanisms (29–31). We also found positions 64 (last *β*B residue) and 74 (small residues on *β*C) strongly interact with pTyr in >80% of structures. As binding transitions to the C-terminal portion, binding mode diversity becomes apparent, again highlighting position 105 as the most C-terminal SH2 residue engaging the pTyr. This visualization also shows ligand position −1 engagement predominantly occurs with adjacent pTyr binding (position 37), while the +1 position is concurrently engaged with position 103, suggesting dual constraints for these ligand residues. However, since other SH2 residues can independently interact with the +1 position (but not −1), there may be differences in constraints between −1 and +1 positions. Comprehensive mapping bridges knowledge from individual structures and highlights domain-specific variations driving different ligand specificities.

We also examined whether our structural analysis could explain phage display mutagenesis results from researchers developing SH2 “superbinders” for pTyr pull-down reagents (7, 23). When mapping phage-based mutation effects (Fig. S8), the mutations increasing binding affinity directly correspond to the N-terminal SH2 domain region coordinating pTyr binding, including positions 74 and 105—two of the triplet mutations yielding 100-fold higher affinity currently used in improved reagents (7) (Fig. 2B). The third mutation in the triplet (position 67) is adjacent to a conserved pTyr interaction residue, suggesting immediately adjacent positions also shape the binding pocket. Non-evolvable positions (losing function when mutated) include the invariant arginine at position 62 and conserved positions not involved in binding—likely affecting domain folding rather than the binding interface. Positions altering SH2 domain specificity occur at and beyond position 124, in the variable structural contact region (Fig. 2B and Fig. S8). Thus, our systematic extraction identifies both conserved structural positions in the pTyr pocket and variable, SH2-specific positions contributing to ligand selection, potentially guiding development of SH2 domain products with altered affinity or specificity.

### The relative proportion of residue-level interactions determines importance of ligand binding

In Figure 2B, we observed that the total number of SH2 domain residues engaged with specific ligand positions correlated with known binding patterns—most interactions occur with the pTyr residue and the C-terminal ligand portion. To examine this further, we extracted the number of residue-level bonds made to each ligand position and normalized by the maximum bonds observed (typically at the pTyr position), Fig. 3. Averaging across all SH2 domains confirmed that most bonds coordinate the pTyr residue, followed by +1 and +3 positions, then +2, with relatively few bonds to the N-terminal side or beyond +3. This pattern aligns with SH2 domain binding profiles from degenerate peptide libraries, where C-terminal positions, particularly +3, are important selection criteria for protein-protein interactions (32, 33).

**Fig. 3.**
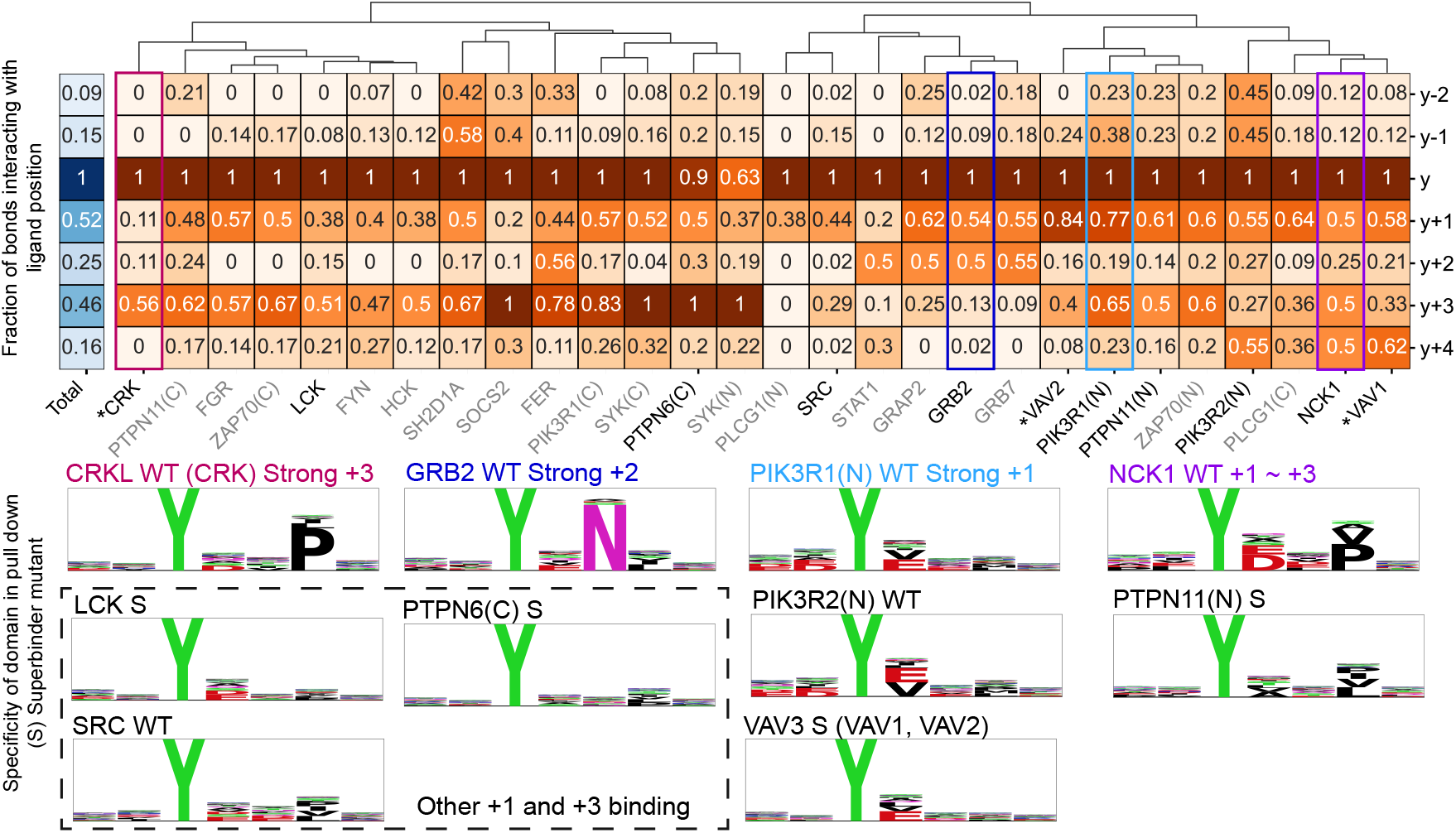
Testing the hypothesis that the fraction of bonds dedicated to coordinating specific ligand positions is related to ligand specificity. For each domain complexed with ligand in available structures, we normalized the number of residue-level interactions made to ligand positions by the maximum number of residue-level bonds made (almost always a value of 1 for the pTyr interaction). ‘Total’ indicates the average across all available structures. The matrix was clustered based on fraction of bonds between 0 and +4 positions. Bottom half of the figure are motif logos generated for peptides pulled down from Martyn et al.’s superbinder experiments of stimulated and pervanadate treated Jurkat cell lysates (7). Here, we included any domain with more than 50 peptides and randomly downsampled to 100 total for controlling entropy and WT indicates the pull down is from wild type SH2 domains and S indicates data is from a “superbinder” mutant domain. Examples are highlighted for the emphasis proportion bonds suggest for selectivity (e.g. strong +3 where the largest fraction occurs with the +3 position and the GRB2 cluster having the highest proportion of +2 interactions). VAV3 and CRKL are not directly represented by available structures, but have close homologs (VAV1/VAV2 and CRK, respectively).

We hypothesized that the fraction of bonds used to coordinate specific ligand positions predicts interaction selectivity. To test this, we analyzed data from a recent study performing pTyr pull-downs from pervanadate-treated Jurkat cell lysates with SH2 domains, including superbinder mutants (7). Figure 3 includes motif logos for domains meeting our information criteria and having ligand-containing structures. While SH2 domains generally engage positions 0 to +3 with emphasis on +1 and +3, there is considerable diversity among family members, revealed by hierarchical clustering of bond fractions. For example, though the average fraction of bonds at the +2 position is relatively low, some members (GRB2, GRAP2) show significantly higher engagement at this position. The GRB2 binding motif is the only one with a significant +2 determinant, consistent with our classification as a “+2 binding mode” group. Additionally, GRB2-related members show relatively low +3 position interactions, consistent with previous findings that a bulky tryptophan in the EF-loop restricts access to the +3 binding pocket (34). Thus, residue-level bond patterns appear predictive of ligand selectivity information content.

Additional binding patterns emerge from the clustered heatmap, including groups with equivalent emphasis on +1 and +3 positions and groups where +1 is stronger than +3. Motif logos generally match these bond fraction observations, suggesting that bond fraction can classify ligand binding modes more broadly. For example, PIK3R1(N) and PIK3R2(N), with stronger +1 bond emphasis, show higher selectivity for +1 than +3 in their motif logos. These domains also have the highest bond fractions at −1 and −2 positions and their logos show the strongest information content in positions N-terminal to pTyr.

While we observe high bond fractions at the +4 position for some domains, no motif logos suggest +4 position selectivity. Pascal et al. found that PLCG1(C), along with other Group II binders (SH2 domains that have small amino acids in the *β*-D5, our alignment position 104), allow for an extended interaction with the ligand (up to and past the +4 ligand position) (35). The lack of ligand discrimination at the pull-down level, despite non-negligible bonds at the +4 position, is likely explained by earlier observations, that extended peptide length does not map new SH2 domain residues – meaning that the +4 interactions are interacting with SH2 domain residues are already interacting with +1 to +3 residues (Figure 2B). Bond fractions for PLCG1(C) suggest strong +1 importance, consistent with mutagenesis studies showing only the +1 position provides significant binding energy outside the pTyr interaction (35). Together, these results demonstrate that comprehensive analysis across structures reveals key information about SH2 domain-ligand coordination and domain-specific binding differences, while highlighting the need for deeper understanding of how combinatorial residue-level interactions determine specificity, as simple bond interaction analysis is insufficient to describe ligand discrimination.

The motif logos indicate, though weakly, preferences for acidic amino acids at the +1 position, consistent with the structural co-dependence between +1 and pTyr binding through SH2 residue 103 (Fig. 2B). Since the content of motif logos assumes an equal distribution of amino acids, we wished to statistically test the presence of a +1 acidic amino acid in the peptides of an individual SH2 domain, compared to the overall background of all peptides identified in the experiment, controlling for sequence aspects related to the condition and ability to be phosphorylated by kinases (Table S1). Strikingly, we saw significant enrichment for E/D in the +1 position for 10 of the 12 total domains there was data for (8 of 10 of the domains for which we also have structures). For example, Nck1 had an FDR corrected p-value of 4.9e-21 and VAV3 had 2.25e-8. The two domains lacking +1 acidic enrichment were CRKL and GRB2, which show the strongest +3 determinants. Testing for −1 acidic enrichment showed CRKL and GRB2 were instead enriched for this constraint, with fewer domains showing −1 enrichment overall and weaker effect sizes. Interestingly, structures capturing the −1/pTyr dual constraint are less common than those capturing pTyr/+1 engagement, correlating with the observed effect sizes in phosphopeptide pulldowns. These independent, high-throughput data suggest our structural analysis is highly informative at both global and individual family levels

### Intraprotein, domain-domain, interaction interfaces

To understand the role of PTMs, mutations, and non-ligand interfaces in regulating full protein architectures, we examined interfaces between SH2 domains and other modular domains within a protein (intraprotein interactions). This analysis allowed us to test domain-domain contact similarity between experimental and AlphaFold structures and assess interface conservation when domain pairs are reused in different protein architectures. PDB structures provided contact maps for 9 domain-domain interfaces spanning 21 unique SH2 domains (Fig S9), while AlphaFold predictions yielded 5 additional interfaces plus comparisons with the 9 from PDB (Fig S10), covering 43 unique SH2 domains and their adjoining domains.

To evaluate AlphaFold predicted structures, we compared domain-domain contact maps from experimental and predicted structures using Jaccard Index (JI)—the number of shared residue features normalized by the total features from both methods. Experimental and predicted contacts showed high similarity (JI >0.5) for 16 of 19 domain-domain interfaces (Fig S11A). For interfaces with low similarity, we examined the Predicted Aligned Error (PAE) from AlphaFold, which estimates distance error between residue pairs. The SH2-PE/DAG-bd interface in CHN2 shared only half the contacts between experiment and prediction and had high average PAE, while the PTPN6 SH2-SH2 interface showed substantial differences with poor PAE values (Data File S2). These results indicate PAE is useful for assessing predicted structure reliability for contact extraction (Fig S11). Based on this analysis, we used predicted structure interfaces only when PAE estimates showed at least 50% of the domain-domain interface within acceptable error range (PAE *≤* 10Å). Given the generally high agreement between experimental and predicted structures, with PAE-based filtering to reduce errors, we generated contact maps for an additional 13 unique interfaces using AlphaFold predictions.

Having validated our approach for both experimental and predicted domain-domain interface extraction, we examined interface composition across the SH2 family and similarity between proteins sharing domain-domain pairings. We found the extent of SH2 domain engagement in domain-domain interfaces varies considerably. For example, the SH2-SOCS_box interface is extensive (41 contacts) in SOCS proteins, while others consist of just a few contacts, such as the tandem SH2 domain interface in the PTP family (Fig. 4A). Interface extent may indicate overall protein regulation importance. For instance, the extensive contacts between SH2(N) and PTP catalytic domains in PTPN6/11 align with the SH2 domain’s role in maintaining phosphatase inactive conformation (36).

**Fig. 4.**
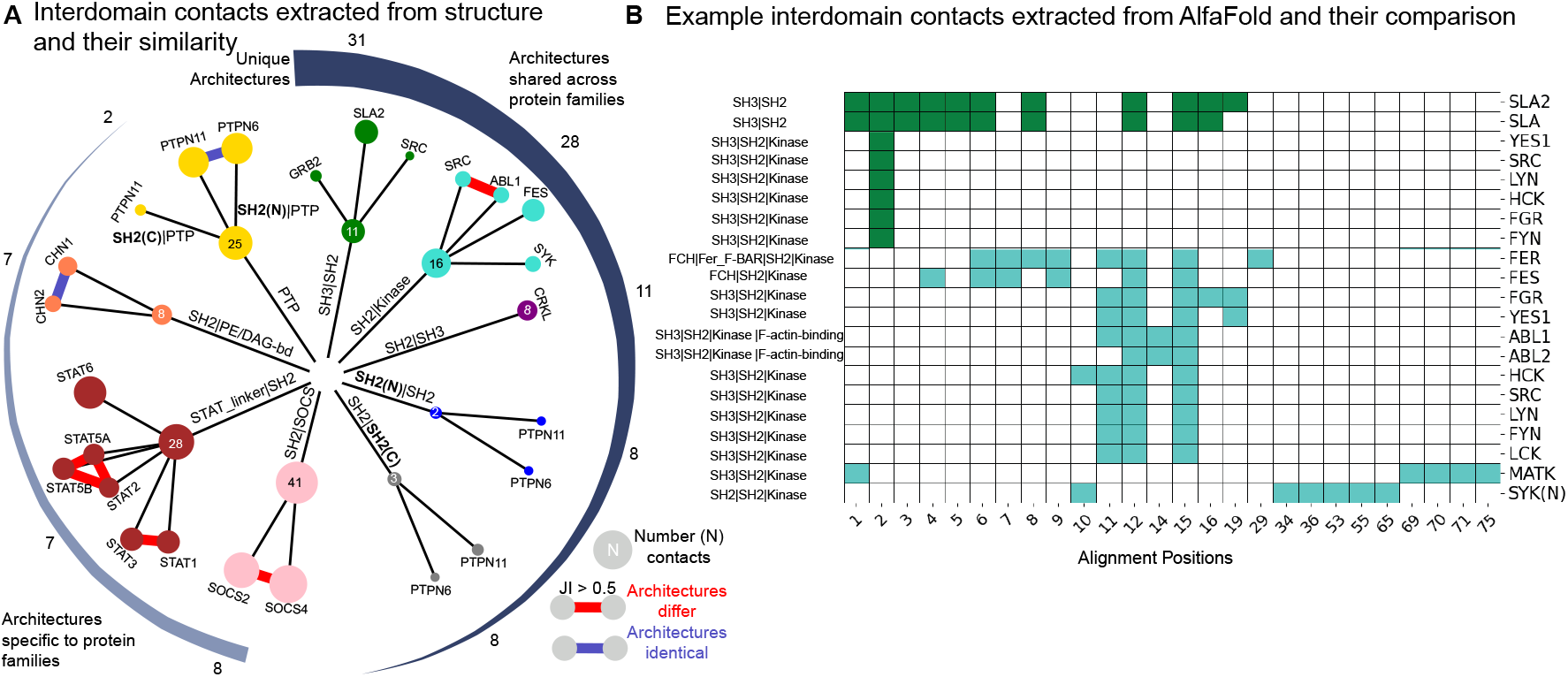
**A)** All domain-domain interactions extracted from available structures. The size of the nodes are proportional to the total number of contacts made with that domain interface and the SH2 domain. If the Jaccard index (JI) of the overlap between features was greater than 0.5 for multiple proteins, the line connects those proteins (red if the full protein architectures are identical, blue if the full protein architectures are different). The interfaces are sorted, in a circular format, according to the size of the total number in the proteome (e.g. 31 SH2 domain-containing proteins have an SH3-SH2 pairing). **B)** AlphaFold predictions for the interfaces of SH3-SH2 (dark green) and SH2-Kinase interfaces (cyan) across diverse full protein architectures. Protein architecture is given on the left and protein name on right. Positions on x-axis are relative to the reference SH2 domain alignment.

When measuring similarity of domain-domain contacts across the family, we found high conservation of interfaces among proteins sharing entire protein architectures (Fig. 4A). Examples include high interface similarity across homologs with identical architectures: PTPN11/PTPN6, CHN1/2, and STAT5A/B. This conservation extends to families with minor architectural variations (e.g., SOCS2/4, JAKs, GRB7/10/14, and STAT1/2/3; Fig. S9 and Fig. S10). However, when domain-domain pairings (sub-architectures) occur across highly variable protein compositions, their interfaces differ substantially. For example, SH2-SH3 interfaces within SLA/SLA2 and SRC family kinases remain conserved only within their respective families (Fig. 4B). The SH2-Kinase interface contains a core of approximately 4-6 amino acids near the SH2 domain’s N-terminus that shows high similarity between ABL and SRC family kinases. This conservation suggests the ABL family’s additional F-actin-binding domain has minimal impact on SH2-kinase domain interactions compared to SRC family kinases. Neither ABL nor SRC families share the same level of SH2-Kinase interface conservation with FER/FES or SYK families (Fig. 4B). Thus, domain-domain interfaces tend to remain conserved among homologous protein duplications and can withstand small architectural expansions but rarely survive major alterations to overall protein architecture.

Given that domain additions can alter structural arrangements of shared sub-architectures, we investigated how contacts depend on experimental design, particularly protein length. We found dramatic differences when comparing partial versus full protein structures. For example, PTPN11, which contains two SH2 domains followed by a tyrosine phosphatase catalytic domain (SH2-SH2-PTP_cat), shows consistent interface contacts when the full protein is expressed. However, structures containing only the tandem SH2 domains show interfaces that differ significantly from those in full-length structures (Fig S12). Conversely, some STAT and SOCS partial protein structures maintain similar interfaces to full structures (confirmed by high-confidence AlphaFold predictions). Since partial protein structures can misrepresent native domain-domain interfaces, we increased our stringency by prioritizing full domain architectures and using partial structures only when domain-domain pairs align well structurally (low RMSD) with full protein structures (Data File S2B). When full structures were unavailable by experimentally-derived structures, we used AlphaFold predictions with acceptable PAE values. This approach produced a reliable set of contact maps between SH2 domains and other protein domains across the family, providing insights into how these domains regulate protein function and folding (Fig. 1B).

### Extending contact maps by evolution and structural similarity

Despite numerous SH2 domain structures, coverage of the family remains limited—only *≈*24% and *≈*18% of SH2 domains have experimental structures with ligands or multiple domains, respectively. We therefore developed a method to confidently group SH2 domains for projecting contact maps from available structures to those lacking structural data, particularly challenging for ligand binding pockets with variable specificity regions. Using evolutionary distances, we hierarchically grouped SH2 domains through a NeighborJoining method (Fig S13), identifying 26 initial clusters. Further analysis revealed the need for sub-clustering; for example, a cluster containing PIK3R(C) and VAV family domains showed strong within-group sequence homology but poor cross-group similarity (Fig S14). Since these cases couldn’t be systematically determined through hierarchical tree cutting alone, we performed clustering within the 26 clusters, resulting in 58 sub-clusters (Data File S1). We validated this approach by comparing with sequence-identity-based and structure-based clustering methods (minimizing RMSD scores). Sequence-based clustering produced identical results, while structure-based clustering reproduced 79% of the clusters with more than two proteins. We used identical sub-clusters found across all approaches, grouping remaining domains together (Data File S1). To test these groupings for ligand contact projection, we measured contact similarity within groups having multiple structures. Generally, we found good agreement, with (GRB2, GRAP2), (VAV1, VAV2), and (HCK, LCK, FGR) groups showing high contact similarity (JI values >0.5, Fig S15). However, tandem SH2 domains revealed a discrepancy: while clustering suggested between-protein domains were more similar (e.g., PTPN6(N) more related to PTPN11(N) than to PTPN6(C)), structural contacts showed within-protein domains had similar binding patterns. This held true across all tandem domains with structural data (SYK, ZAP70, PTPN11, PIK3R1). We therefore handled tandem SH2 domains as separate groups (Data File S1C), yielding 42 final clusters for ligand contact projection (Data File S1). Based on our finding that domain-domain interfaces are most conserved when full protein architectures are shared, we used protein architecture for domain-domain contact projection. Using projections from available structures within clusters for ligand contacts, and from clusters with identical architectures for domain-domain contacts, we extended contact maps to 31 (ligand) and 10 (domain-domain) additional SH2 domains (Fig. 1B). While relaxing constraints could increase coverage, we prioritized high structural and evolutionary similarity for reliable projections.

### Inferring the effect of post-translational modifications by conserved structural analysis

SH2 domains undergo extensive post-translational modification. The 119 human SH2 domains contain 191 pTyr sites, 164 ubiquitinated lysines, 168 phosphorylated serines, 65 phosphorylated threonines, 34 acetylated lysines, and smaller numbers of methylations and sumoylations that have been identified experimentally. Despite these numerous modifications, few studies report their functional effects. Notable exceptions include independent reports on SRC family kinases where SH2 domain phosphorylation alters binding. In LCK, Y192 phosphorylation decreases pY ligand binding affinity, particularly affecting the interaction with the ligand’s +3 position (12), reducing LCK activity in TCR signaling (37). Similarly, phosphorylation of LYN Y194 decreases peptide binding regardless of the peptide’s inherent affinity (13). SRC Y213 phosphorylation reduces binding to its C-terminal tail phosphotyrosine (Y527) but not to other ligands (11). These phosphotyrosines occupy the same structural position (alignment position 124), adjacent to a conserved +3 binding site at position 125 (Fig. 2A). Their location in the specificity-determining region near +3 binding residues explains how SRC family kinase SH2 domains maintain binding capacity but alter specificity when phosphorylated at this position, particularly affecting +3 determinants. Given the general importance of the +3 position, this modification can appear to cause substantial binding reduction, as observed for LYN Y194 (13). We hypothesized that integrating PTMs with comprehensive structural analysis might enable faster functional prediction for the numerous SH2 domain PTMs.

We used CoDIAC modules with ProteomeScout (3), PhosphoSitePlus (19), and Jalview to map PTMs onto SH2 reference sequences and analyze them in the PROMALS3D alignment. This revealed patterns of conserved PTMs across structural positions, with multiple modifications appearing at specific alignment positions across many family members. We developed comprehensive reports analyzing each PTM’s relationship to the number of similar PTMs at the same structural position, distance to ligand binding interfaces, nearby interface residues, and overlap with domain-domain or phospholipid binding interfaces (Data File S3).

From this analysis, we generated hypotheses about PTM functional impacts. Based on size and charge differences, we propose that modifications at or near the pTyr binding interface may block domain-ligand binding entirely, while those farther from pTyr likely modulate specificity or affinity. We identified 53 modifications directly on pTyr-engaging residues that could disrupt canonical binding: 18 pY, 29 pS/pT, and 7 N6-acetyllysine sites. Despite limited overall acetylation data, several modifications occur directly at interaction interfaces, notably on a highly conserved lysine at position 105 involved in pTyr coordination (plus 5 additional acetylations on specificity region residues). This suggests acetylation may directly disrupt pTyr-mediated signaling by removing positive charges critical for pTyr coordination.

Phosphorylation shows distinct patterns. Conserved phospho-serine/threonine sites predominate in the N-terminal SH2 domain half, frequently on residues directly contacting the ligand pTyr, suggesting pSer/pThr signaling might directly dampen SH2-pTyr interactions. In contrast, tyrosine phosphorylation tends to occur on or near residues binding the ligand +1 to +3 region, suggesting pTyr might “tune” binding specificity. However, the most conserved tyrosine phosphorylation site (position 73, Fig. 5) sits adjacent to a structural position binding the ligand pTyr on the *β*-C strand, indicating some pTyr sites may function as binding switches rather than specificity modulators.

**Fig. 5.**
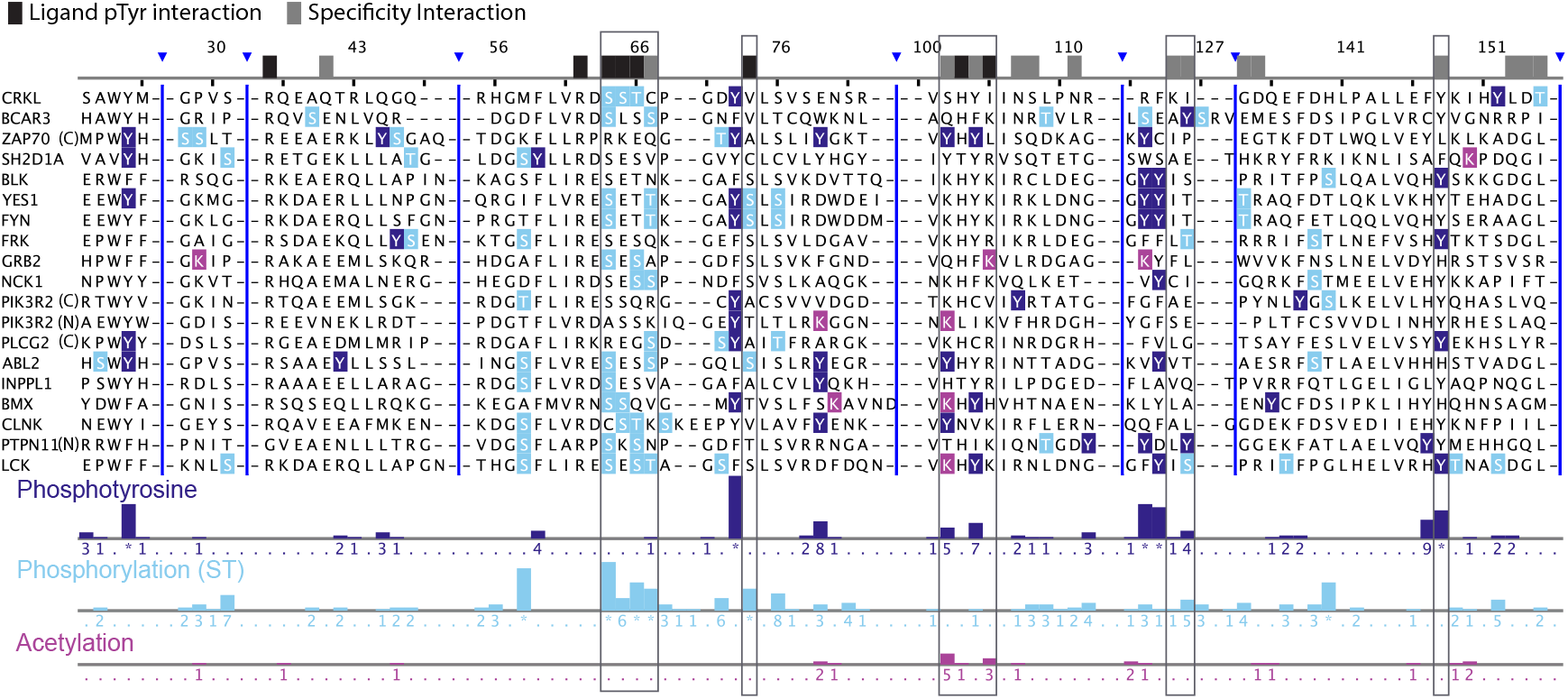
Post-translational modifications (PTMs) in a Jalview-based visualization on the reference SH2 domain alignment for a subset of SH2 domains that cover the diversity of PTMs. Ligand features were collapsed across structures and indicated if they were shared by 10 or more unique SH2-ligand pairs. Ligand interactions were labeled as interacting with pTyr (black) or specificity (gray) based on whether the interaction was predominantly with the pTyr residue or with other ligand positions (−1 to +3 predominantly). Tracks at the bottom indicate number of PTMs found in that alignment position across the family. Numbers on the tracks indicate the number of PTMs identified in that position, * indicates more than 10 (pTyr: position 73 has 28, position 123, 124, and 148 have 16, 14, 13, respectively; pSer/pThr position 58 has 19, and the positions between 64 and 67 have 22, 6, 13, and 10 phosphorylation sites, 74 has 10 pSer/pThr sites. Dark blue indicates phosphotyrosine; Light blue indicates a phospho-serine/threonine; Maroon indicates N6-acetyllysine. Boxes around alignment and tracks indicate locations of high conservation of PTMs and overlap with ligand interacting residues.

Given the extensive phosphorylation at or near ligand binding residues, we searched for functional studies of these modifications. Literature confirms that some pSer/pThr (38) and pTyr sites (39) affect binding and signaling. For example, Lee et al. studied two phosphoserine sites in PIK3R1 (p85a) in the same structural position of both tandem SH2 domains (alignment position 65, a conserved pSer/pThr site). Phosphorylation of either site dramatically decreased SH2 domain ligand binding and inhibited PI3K signaling (38). When both sites were phosphorylated, PI3K dissociated from upstream activators, reducing AKT activation. This region of the SH2 domain contains 51 annotated phosphoserine/threonine sites directly interacting with ligands, suggesting similar regulatory effects across many SH2 domains.

For tyrosine phosphorylation, our conservation analysis identified the previously studied SRC family kinase regulatory sites and extended them beyond SFK SH2 domains—alignment position 124 contains 14 modification sites including in ZAP70(C), NCK1, and ABL2 (Data File S3) SH2 domains. This position neighbors conserved specificity contacts at positions 125-126, consistent with phosphorylation affecting +2/+3 interactions (Fig. 2B). We also observed many SH2 domains contain phosphorylated tyrosine pairs (positions 123-124), including in SRC family kinases. Weir et al. found three tyrosines in FYN (Y185, Y213, Y214; alignment positions 73, 123, 124—all conserved phosphorylation sites) reduced SH2 domain binding capacity when phosphorylated (39). These studies validate our comprehensive contact mapping approach for predicting PTM effects. We estimate 54% and 35% of SH2 domains can be regulated by tyrosine and serine/threonine phosphorylation respectively, often with multiple regulatory sites on a single domain.

Beyond ligand binding, we examined PTMs at domain-domain interaction interfaces to identify potential regulation of larger protein structures. Due to the architecture-specific nature of these interfaces, we focused on PTMs directly at domain-domain interfaces, identifying 40 such modifications (Data File S3). This approach revealed key regulatory insights, including Burmeister et al.’s finding that phosphorylation of PTPN11(N) T73 and PTPN11(C) S189 (alignment position 140) by PKA inhibits PTPN11 catalytic activity (40) by stabilizing its closed conformation. Our analysis confirms PTPN11(N) T73 is at the SH2-PTP catalytic domain interface and PTPN11(C) S189 at the SH2-SH2 domain interface, supporting their role in regulating protein conformation. In total, we identified 28 PTMs at SH2-domain interfaces with other domains (Data File S3), providing directed hypotheses for functional testing.

We also examined phospholipid-binding interfaces using hand-annotated features from Park et al. (24). With data for about 10 SH2 domains having phospholipid binding residues, we identified five pTyr sites across four domains that might affect lipid binding through proximity to positively charged interface residues (Data File S3). Interestingly, several phospholipid-binding lysines (6 total) have been annotated as ubiquitination sites, suggesting potential regulation of phospholipid binding, though surface accessibility may also explain this pattern.

### Integrated analysis of clinically relevant mutations

We used CoDIAC to connect mutations to SH2 domain regulation by examining relationships between mutations, interaction interfaces, and PTMs (Data File S4). From 111 clinically significant mutations identified across databases (OMIM, gnomAD, PDB), we found 29 on ligand contacts, 40 within 2 amino acids of ligand binding, 16 at domain-domain contacts, and 9 on modified residues. Across ligand and domain-domain contacts, 83-85% of mutations drastically change residue physiochemical properties, compared to 44% at PTM sites (Data File S4).

Integrated analysis revealed diverse potential effects. For example, SRC K206L (position 105) removes a positive charge at a pTyr-coordinating residue; STAT1 K637E (position 134) introduces a charge switch at a ligandbinding residue and eliminates a ubiquitination site; and PTPN11 Y62D (position 112) affects a residue adjacent to the ligand interface and at the PTP catalytic interface, while removing a phosphorylation site. PTPN11 Y62D and Y63 mutations have been associated with Noonan syndrome, Leopard syndrome, and RASopathy (ClinVar accessions: VCV000013329.48, VCV000013333.86). Mutations disrupting the SH2-PTP interface increase catalytic activity by destabilizing the closed conformation (41). Our analysis shows these mutations not only affect domain-domain interfaces but may also disrupt ligand binding and phosphorylation-based regulation.

Another example from integrated analysis involves SOCS1 mutations. P123R and Y154H mutations activate the JAK-STAT pathway in tumor cells and are associated with B-cell lymphomas (42). We found P123 (position 101) is directly adjacent to a conserved +1 ligand binding residue (position 102) and on the same *β*-D strand side as the pTyr/+1 interacting residue 103 (Fig. 2B). SOCS1 Y154 (alignment position 148) is both a ligand binding contact and interacts with the SOCS_box domain. While Y154 phosphorylation hasn’t been identified in SOCS1, it aligns with a conserved phosphorylation position predicted to alter binding specificity through +3 ligand interactions. As one of the rare cases where a ligand pocket residue also participates in intraprotein interactions, phosphorylation might regulate SH2 domain opening for ligands. Y154 is part of a “YY” doublet where both tyrosines can be phosphorylated, suggesting the Y154H mutation could affect: 1) SH2-SOCS_box interface disruption, altering catalytic activity; 2) ligand specificity changes; and 3) phosphorylation-based regulation of both ligand and domain-domain interfaces. While previous work hypothesized ligand contact loss (42), our analysis precisely identifies affected interactions and suggests additional regulatory mechanisms.

## Discussion

CoDIAC provides a flexible framework for extracting contact maps from experimental and predicted structures, annotating domains within structures, and harnessing all available structures for protein domain families. Here, we used it to explore interaction interfaces of modular domains, creating a comprehensive overview of SH2 domains and the intersection of post-translational modifications and mutations with binding interfaces. While recovering insights from individual structure-based studies, the comprehensive evaluation across the entire family revealed emergent properties of SH2 domains.

A key limitation of this and all structure-based studies is that interaction interfaces, particularly domain-domain contacts, represent single low-energy configurations and do not capture the dynamic nature of protein interactions. Nevertheless, these low-energy states remain relevant for understanding how mutations and PTMs affect protein configuration. Although we did not examine the role of linker regions in regulating protein function—a well-known phenomenon in SH2 domain-containing tyrosine kinases (43) – CoDIAC can readily handle these regions, as interdomain regions are effectively annotated by the pipeline. Additionally, some structures note domain-domain interfaces that change in active conformational states, such as an SH2-kinase interface in active ABL that stabilizes the open conformation (43). Such interfaces are also important, including the possibility that PTMs could regulate conformational state stability. By selecting contacts shared by the majority of structures to reduce study bias, we inherently biased our maps toward the most represented structures (typically inactive conformations). However, CoDIAC can easily compare and contrast different interfaces for such comparative purposes and can map non-domain regions and various interface types, including interactions with lipids, small molecules, RNA, and DNA.

Our study highlighted several important caveats. Structures with partial protein coverage should be used cautiously, as demonstrated by the differences in contact maps between partial and full protein representations and the structural rearrangements that can occur when sub-architectures are shared between diverse families. Additionally, we observed that ligand engagement can occur without the canonical pTyr-invariant arginine interaction when ligands are presented in cis or through multivalent interactions (as in tandem SH2 domains). This suggests contact mapping from isolated domain-ligand pairs may limit our understanding of physiologically relevant interactions in signaling networks. Beyond tandem SH2 domains, many domain-ligand interacting modules work together with other modules (e.g., the SH3-SH2 pairing across much of the family), suggesting that non-canonical binding might be more prevalent in protein interaction networks than isolated domain studies would indicate.

Conservation analysis of PTMs within SH2 domains revealed patterns suggesting extensive regulation of SH2 domain interaction interfaces by multiple signaling systems. Some insights were surprising, contradicting the prevailing notion that modifications primarily occur in intrinsically disordered segments – many conserved sites are located directly on *β*-strands within the SH2 domain. The breadth of these PTMs across independent experiments and homologs, along with evidence of their regulation by drugs (44), growth factors (45, 46), or other stimuli (47), strongly suggests these structurally conserved modifications are transiently regulated, important for signaling, and not simply mass spectrometry artifacts. Literature describing modification effects in conserved positions on sites not yet in databases also suggests additional modifiable residues in these positions may be discovered.

While we highlighted individual PTM effects on regulating ligand or domain-domain interactions, evidence suggests they co-occur and provide multifactorial control of SH2 domains. An intriguing pattern involves phosphoserine sites directly adjacent to phosphotyrosine sites near key ligand binding regions, including positions 72 and 73 which interact with the ligand pTyr. Recent profiling of human serine/threonine kinases revealed that priming phosphorylation may be common for subsequent phosphorylation (48). Using ProteomeScout, we found evidence of a doubly phosphorylated tryptic fragment containing these positions in the SH2D1A domain from breast cancer samples (49), suggesting one phosphorylation may prime the kinase motif for the other. While much remains to be tested about PTM regulation of SH2 domain interactions, the comprehensive integration of structure and PTMs enables more direct hypothesis generation about modification effects and prioritization based on conservation and functional impact, while identifying multiple factors to consider in disease-relevant mutations.

## Materials and Methods

### UniProt, InterPro, and structure reference generation of the SH2 domain family

We developed and used the CoDIAC ‘InterPro’ module to gather all SH2 domain containing proteins for ‘Homo Sapiens’ with the InterPro ID ‘IPR000980’, resulting in 109 UniProt identifiers. We then used the ‘UniProt’ module of CoDIAC to build reference files of the proteins and a reference FASTA file of just the SH2 domain regions for the 119 unique SH2 domains. Given that there can sometimes be discrepancies around specific domain boundaries, we included the ability to alter the boundaries (systematically for all domains) when producing a FASTA sequence file of domains. Based on alignment quality, we found that truncating the boundary of the SH2 domains defined in the reference by one amino acid on the C-terminal resulted in a significantly better alignment and we selected an ‘n-terminal offset’ of 0 and a ‘c-terminal offset’ of −1. Additionally on domain quality checks, given its high specificity for only the PIK3 regulatory family, we manually removed the InterPro region in the PIK3R1/2/3 proteins ‘PI3K_P85_iSH2:IPR032498’, which is an alpha-helical region of between the tandem SH2 domains. We also manually removed an InterPro defined domain in SUPT6H that indicates (Spt6_SH2_C:IPR035018) as it overlapped with the parent SH2 family. Finally, we found that using the SMART (50) defined boundaries, which were an average between UniProt and InterPro domains for the atypical SH2 domains (JAK family) resulted in a better overall alignment. At the time of this analysis (June 2024), UniProt returned 1,477 experimental structures associated with the UniProt set of proteins.

We created the structural reference datasets containing structures for SH2 domain proteins obtained through experiments (Integrated PDB reference file) and predictions (AlphaFold reference file). For the generation of this dataset, we used CoDIAC’s ‘PDB’ followed by ‘IntegrateStructure_Reference’ module that captured an exhaustive list of experimental details for each of the PDB identifiers associated to the UniProt IDs of human proteins containing an SH2 domain. Using CoDIAC’s ‘IntegrateStructure’ module, which aligns structure data with the reference data and annotates domains found within structures, we found 467 total structures for analysis that covered an SH2 domain in its entirety. Gaps, and variants of experimentally-derived sequences, relative to reference, were noted in the annotated file for consideration of exclusion criteria during contact mapping. We performed an identical process of capturing all predicted structures for the UniProt IDs identified in the family, annotating their regions relative to the reference and generated the AlphaFold reference file. The 109 predicted structures were downloaded as mmCIF files using AlphaFold database version 2 (v2). We used CoDIAC’s ‘PTM’ module to capture PTMs found within the domain boundary regions of SH2 domains for all PTMs for which there were five or more PTMs in the family, producing a unique set of PTMs from both ProteomeScout (3), using the ProteomeScoutAPI (51), and PhosphoSitePlus (19) using an API that we created and incorporated in CoDIAC that operates similarly to the ProteomeScoutAPI, then combined them and kept the unique set of all PTMs for final analysis. We used CoDIAC’s ‘mutations’ module to gather mutations within the SH2 domain regions defined in the reference from gnomAD and OMIM. We used PROMALS3D (26) to align the reference SH2 FASTA file and then used CoDIAC’s ‘Jalview’ module to translate and integrate feature files produced from PTMs along with contact map feature files, along with creating annotation tracks of features based on summing features along the columns of the alignment. Integrated features and annotation tracks were used to create analysis reports concerning PTMs and mutations (compiled in data S3 and S4, respectively). All data generated by CoDIAC version 1.0.0 are available at https://doi.org/10.6084/m9.figshare.26321968 as well as on our GitHub repository, where we also have provided all Python code we developed for SH2 domain analysis at https://github.com/NaegleLab/SH2_contact_analysis.

### Adjacency File and Contact Map generation

We generated binary text files using the ‘AdjacencyFiles’ module that are a simplified representation of interatomic interactions. To make these files, we first make these calculations using a Python package Arpeggio (25), which outputs a JSON-formatted file comprising all types of interatomic interactions that exists within an input protein structure. For the purposes of domain focused contact extraction, we kept all non-covalent interactions (aromatic, carbonyl, hydrophobic, ionic, polar, van der Waals, halogen, and hydrogen bond) occurring at a distance < 5Å between atoms that reside on SH2 domains and their interacting domains/ligands. We do not include any interactions that may occur between protein entities and small molecules in this work. All components (chains, residues, residue positions, atom pairs, distance, and contact type) of the filtered non-covalent interactions are saved as adjacency text files and using our set threshold (retain interactions if found in >=25% of the chains for domain-domain analysis and in >=50% of bound chain replicates for domain-ligand complexes) we aggregate contacts across chains of specific entities. We represented whether the contact exists (binary value ‘1’) or not (binary value ‘0’) between or across protein entities in the final binarized adjacency text files. We were unable to generate Arpeggio json files for 12 PDB structures (PDB IDs: 7UNC, 7UND, 8H36, 8H37, 8OEU, 8OEV, 8OF0, 6GMH, 6TED, 7OOP, 7OPC, 7OPD) and the adjacency files we successfully generated for 455 PDB and 109 AlphaFold structures generated in this work can be found here (https://doi.org/10.6084/m9.figshare.26309674). Across the 475 PDB structures, we identified 17 unique post translational modifications (PTM). We generated a PTM dictionary that is used by the ‘contactMap’ module to replace these modified residue three letter codes by their native residue’s single-letter abbreviation in the structure sequence and allowed for precise comparisons between structure and reference sequence. This PTM dictionary can be modified based on the ones extracted within the structure datasets.

We used CoDIAC’s ‘contactMap’ module to generate contact maps for domain-domain and ligand binding interfaces of SH2 domains. This module creates a Python object that stores all the structural information gathered from both the structure datasets (PDB/AlphaFold reference files) and the binary adjacency files. By incorporating annotated and interaction data for each structure, contact maps were constructed in the form of nested python dictionaries whose outer and inner dictionary keys indicate the residues that are non-covalently linked and the values for the inner dictionary represents the binary value of the interaction. We printed contacts from these dictionaries to Jalview supported feature files that bind SH2 domains and its other interacting domains/ligands. Using the ‘analysis’ module we aggregated contacts across various PDB structures that represent an identical interface. We evaluated after what threshold (percent) the loss of features upon aggregation becomes minimal and found 30% to be suitable threshold for merging contacts across SH2 domain containing structures (Fig S1).

We found 207 PDB structures spanning 32 unique proteins that allowed us to examine domain-domain interfaces for contacts. We removed PDB structures for LCK, PLCG1, ZAP70 from our analysis since we found structural discrepancies while evaluating RMSDs and were unsure whether the structures represented a native conformation. We do not extract contacts across SH2 domains and other modular domains within PIK3R1, CRK and SUPT6H structures. Given these removals and the evaluation of wild-type structures for domain-domain binding interfaces resulted in maps for 19 unique SH2 domain proteins (Fig. 1B). We failed to generate contact maps due to a mismatch between the structure and reference sequence alignments for two of the HCK structures (PDB IDs: 2HCK and 1AD5), due to a 10 amino acid unmodeled region within their kinase domains. For the domain-ligand contact analysis, we excluded SH2-pTyr complex structures (PDB IDs: 1A07, 1BF5, 1A81, 1CSY, 1CSZ, 2DVJ, 2LQW, 2OQ1, 4Y5U, 4Y5W, 4XZ1, 5D39, 7Q5U, 7Q5W, 7Q5T) identified as binding in a non-canonical manner (i.e., not involving the invariant ‘R’ interaction).

### PAE estimation for predicted structural fold of domain-domain interfaces

AlphaFold prediction system provides Predicted Aligned Error (PAE) values for every residue pair in a protein structure that reflects the distance error (Å) between the residues oriented in three dimensional space. Given a domain interface of interest, we first retrieve the PAE values between the residue pairs *r*_*i*_ *r*_*j*_, where *r*_*i*_ and *r*_*j*_ reside on either of the domains. Since AlphaFold presents the PAE values in the form of an asymmetrical matrix, we calculate the average of the PAE estimates for the two relative structural configurations of the domain-domain interface (i.e. from both directions of the interface).

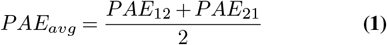

Where *PAE*_12_ is the fraction of total number of residue pairs (*r*_*i*_ − *r*_*j*_) that have a low PAE (<= 10Å), given *r*_*i*_ is on the SH2 domain and *r*_*j*_ on the partner domain and *PAE*_21_ is the estimate of the same interface when *r*_*i*_ is on the partner domain and *r*_*j*_ is the SH2 domain.

We report the final PAE estimate as this average value calculated from both directions of the domain-domain’s structural orientation. We consider assessing interfaces only if our PAE estimate results in capturing at least 50% of the domain-domain interface within our defined tolerable error range.

### Methods for measuring the evolutionary and structural properties of SH2-domains

First, we used evolutionary distances as the basis to classify SH2-domain containing proteins into families. We implemented a distance-based Neighbor-Joining algorithm to calculate the evolutionary distances among all the known 119 SH2 domains. We then applied Ward’s method on the distance matrix to hierarchically cluster the proteins. Using the elbow method identified an optimal linkage distance for clustering – elbow point occurring at 0.5 indicates a decrease in variance within the clusters beyond this linkage distance (Fig S16). This threshold produced 40 clusters (Data File S1D), but we were unable to sufficiently project contacts, so we reassessed clustering for this purpose at the next elbow point that occurs at linkage distance 0.8 that produced 26 clusters (Data File S1A). The clusters generated from a threshold of 0.5 and 0.8 were highly similar. Each of these 26 clusters were subdivided into 58 sub-clusters and we used silhouette score to identify the number of sub-clusters that can be formed within each of these clusters. We benefited by working with a higher linkage threshold of 0.8, since it did not affect the protein grouping and allowed us to find more closely related neighbors for enhancing the chances of contact projection. We gained features for 9 additional proteins at a threshold 0.8, compared to what we would achieve with 0.5. Upon performing these steps, we were able to split the SH2 domain family into 26 clusters and 58 sub-clusters (Data File S1B). We next validated clusters by closely examining the sequence and structure of the domains to evaluate whether the evolutionary-based agglomerative clustering would be beneficial in identifying and separating intra-clustered homologues efficiently. For primary sequence comparison we calculated amino acid sequence identities between proteins within each of these 26 clusters. We hierarchically grouped each of the 26 clusters based on the sequence identity scores. This resulted in exactly the same sub-grouping we had attained using evolutionary distances.

For structural comparisons, we estimate RMSD values using a PyMOL utility ‘super’ that involves sequence-independent programming alignment. We make these calculations between several structures by using Python controlled PyMOL scripts. The experimental structures do not cover the entire SH2 domain family, so we covered for the unresolved protein domains by incorporating the predicted structures of the domains from AlphaFold, since we observed very low RMSD scores (<2Å) between the SH2 domain structures from experiments and AlphaFold (Fig S17). We next repeated similar hierarchical clustering method for the 26 clusters to gain the intra-clustered structural homologues using the RMSD scores.

### pTyr ligand peptide clustering

We represented every pTyr peptide in our dataset as DPPS vectors (52) and then measured the average Euclidean distance between these DPPS vectors of the peptides. These average Euclidean distances indicate how closely associated the peptides are by their physiochemical properties. We hierarchically clustered the peptides based on these average distances. We adapted this method in our work from (8) to categorize pTyr peptides of length 7 amino acids (−2 to +4 of pTyr).

### Phosphopeptide sequence analysis from Martyn et al

Orthogonal testing of structure-extracted features was performed by aligning the sequences phopshopeptide pull downs from pervanadate treated Jurkat cell lysates (7). To generate motif logos, we required at least 50 sequences. If a pull down resulted in significantly more peptides for any one SH2 domains, we subsampled to 100 peptides to control for entropy differences (we did not see differences on the general motif logos across random subsamples). When available, we used the wild type SH2 domain structure. However, for those domains where the wild type structure recovered too few peptides for useful motif generation, we used data from the triplet superbinder mutations. These positions were shown by the authors not to have a significant effect on binding specificity, which is also soundly in agreement with the identification that these positions are exclusively coordinating the pTyr residue, according to our structure extraction (Fig 2B). For testing the preference of acidic amino acids in the SH2 domain pull downs, we first assembled a background of all unique phosphopeptides observed across the various pull down experiments and then tested for the presence of an E or D, directly following or preceeding the phosphotyrosine and used the Fisher’s Exact test to measure the probability of having observed the incidence of the motif by random chance, given the full background. We corrected for multiple hypothesis correction (within the positional group of tests) using the False Discovery Rate procedure (FDR) and rejected hypotheses with FDR corrected values of less than 0.05.

## ACKNOWLEDGEMENTS

We would like to acknowledge: Margaret Ryan for early work on PDB interfaces for identifying relevant SH2 domain containing proteins; Dr. Eli Draizen for help on initial scripts for parsing and contact extractions using mmCIF files; and Dr. Philip Bourne and Dr. Mohammad Fallahi-Sichani for helpful discussions. Research reported in this publication was supported by the National Institute Of General Medical Sciences of the National Institutes of Health under Award Number R35GM138127 and the National Institute of Allergy and Infectious Disease under Award Number R01AI153617. The content is solely the responsibility of the authors and does not necessarily represent the official views of the National Institutes of Health.

## Supplementary Materials

**Fig. S1**. Estimation of a threshold for merging contacts across various binding interface extracted from PDB structures.

**Fig. S2**. Structural coverage for PDB structures resolved for multi-entity protein complexes (SH2-pTyr)

**Fig. S3**. Structural coverage for PDB structures resolved for multi-domains.

**Fig. S4**. Examples of SH2 domain-ligand interactions that do not include the invariant arginine-pTyr interaction (i.e. non-canonical interactions).

**Fig. S5**. SH2-centric features from across every unique SH2-pTyr peptide pair from PDB structures.

**Fig. S6**. Relationship between the contact maps and physiochemical property based grouping of pTyr peptides.

**Fig. S7**. Specific contacts made by ‘pTyr’ with the residues on SH2 domains

**Fig. S8**. Projection of the superbinder mutations on the reference aligned SH2 sequence

**Fig. S9**. Heatmap of domain-domain contacts obtained from experimental structures.

**Fig. S10**. Heatmap of domain-domain contacts obtained from AlphaFold structures.

**Fig. S11**. Comparison of domain-domain contacts extracted from experimental and predicted structures.

**Fig. S12**. Analysis of domain-domain interfaces for partial versus complete experimental structures.

**Fig. S13**. Dendrogram to illustrate the grouping the 119 human SH2-domains using relative evolutionary distances.

**Fig. S14**. Sequence and structure similarity for each of the classified SH2 domain clusters.

**Fig. S15**. Comparison of experimentally determined ligand contact maps across clustered proteins.

**Fig. S16**. Optimum threshold for SH2 domain protein classification using elbow method.

**Fig. S17**. Structural comparison of experimentally and predicted SH2 domain structures.

**Table S1**. Acidic amino acid enrichment testing in SH2 domain pull down experiments

