## Supplementary Figures for "CoDIAC: A comprehensive approach for interaction analysis reveals novel insights into SH2 domain function and regulation"

### This file includes:

**Supplementary Figure 1**: Estimation of a threshold for merging contacts across various binding interface extracted from PDB structures.
**Supplementary Figure 2**: Structural coverage for PDB structures resolved for multi-entity protein complexes (SH2-pTyr)
**Supplementary Figure 3**: Structural coverage for PDB structures resolved for multi-domains.
**Supplementary Figure 4**: Examples of SH2 domain-ligand interactions that do not include the invariant arginine-pTyr interaction (i.e. non-canonical interactions).
**Supplementary Figure 5**: SH2-centric features for every unique SH2-pTyr peptide pair from PDB structures.
**Supplementary Figure 5**: Relationship between the contact maps and physiochemical property based grouping of pTyr peptides
**Supplementary Figure 6**: Specific contacts made by 'pTyr' with the residues on SH2 domains.
**Supplementary Figure 7**: Projection of the superbinder mutations on the reference aligned SH2 sequence
**Supplementary Figure 8**: Heatmap of domain-domain contacts obtained from experimental structures.
**Supplementary Figure 9**: Heatmap of domain-domain contacts obtained from AlphaFold structures.
**Supplementary Figure 10**: Comparison of domain-domain contacts extracted from experimental and predicted structures.
**Supplementary Figure 11**: Analysis of domain-domain interfaces for partial versus complete experimental structures.
**Supplementary Figure 12**: Dendrogram to illustrate the grouping the 119 human SH2-domains using relative evolutionary distances.
**Supplementary Figure 13**: Sequence and structure similarity for each of the classified SH2 domain clusters.
**Supplementary Figure 14**: Comparison of experimentally determined ligand contact maps across clustered proteins.
**Supplementary Figure 15**: Optimum threshold for SH2 domain protein classification using elbow method.
**Supplementary Figure 16**: Structural comparison of experimentally and predicted SH2 domain structures.
**Supplementary Table 1**: Statistical testing of acidic amino acids in the -1 ([DE]y) and +1 (y[DE]) positions.
**Captions for Data File S1-S4**

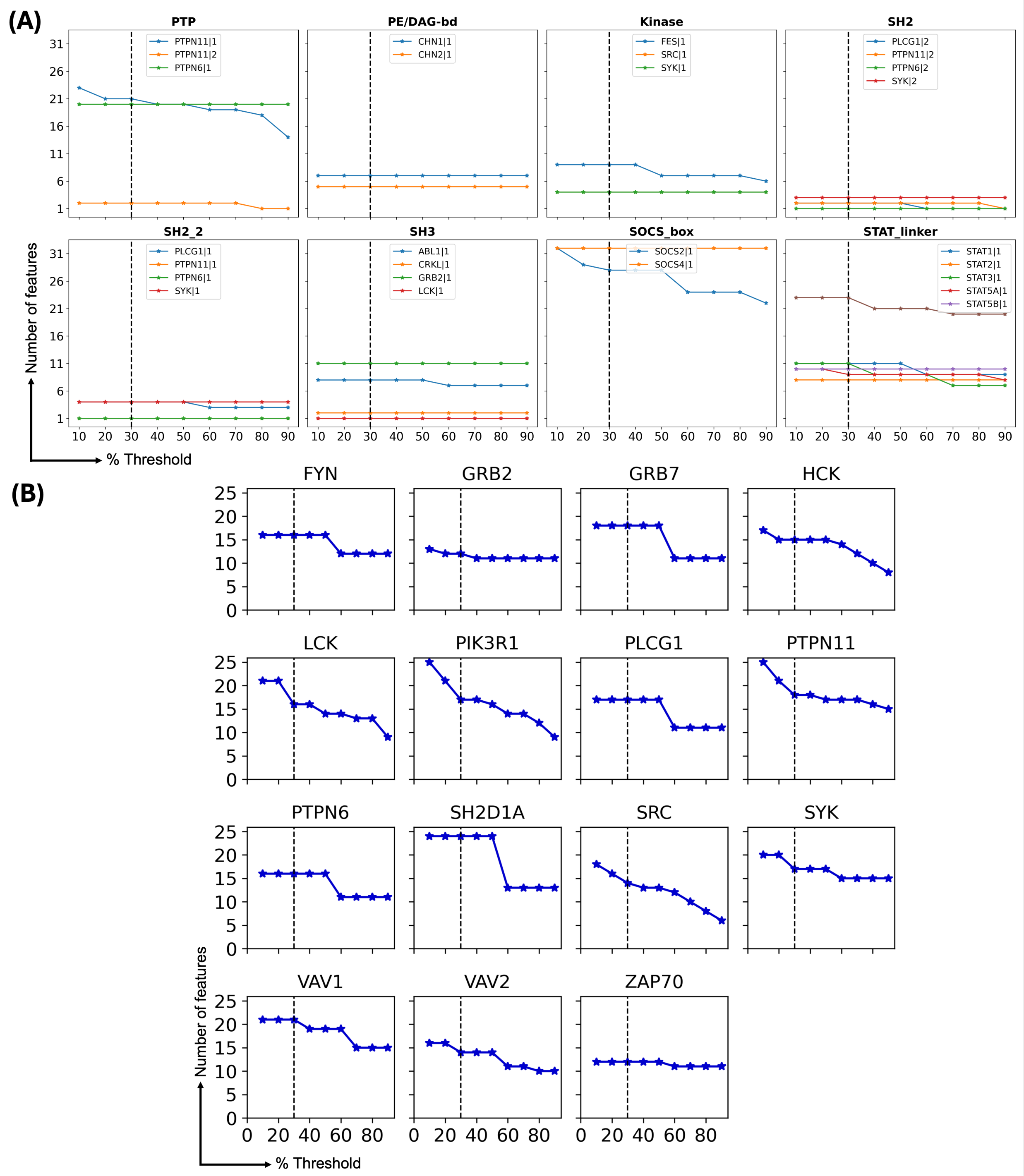


**Supplementary Figure 1. Estimation of a threshold for merging contacts across various binding interfaces extracted from PDB structures.** (**A**) Number of features extracted from domain-domain interfaces based versus threshold (percent of structures that recover a feature). (**B**) Number of features extracted from domain-ligand interfaces versus threshold. The selected threshold (30%) is indicated in the dashed line, where the elbow criterion appears across most decisions for both domain-domain and domain-ligand interfaces.

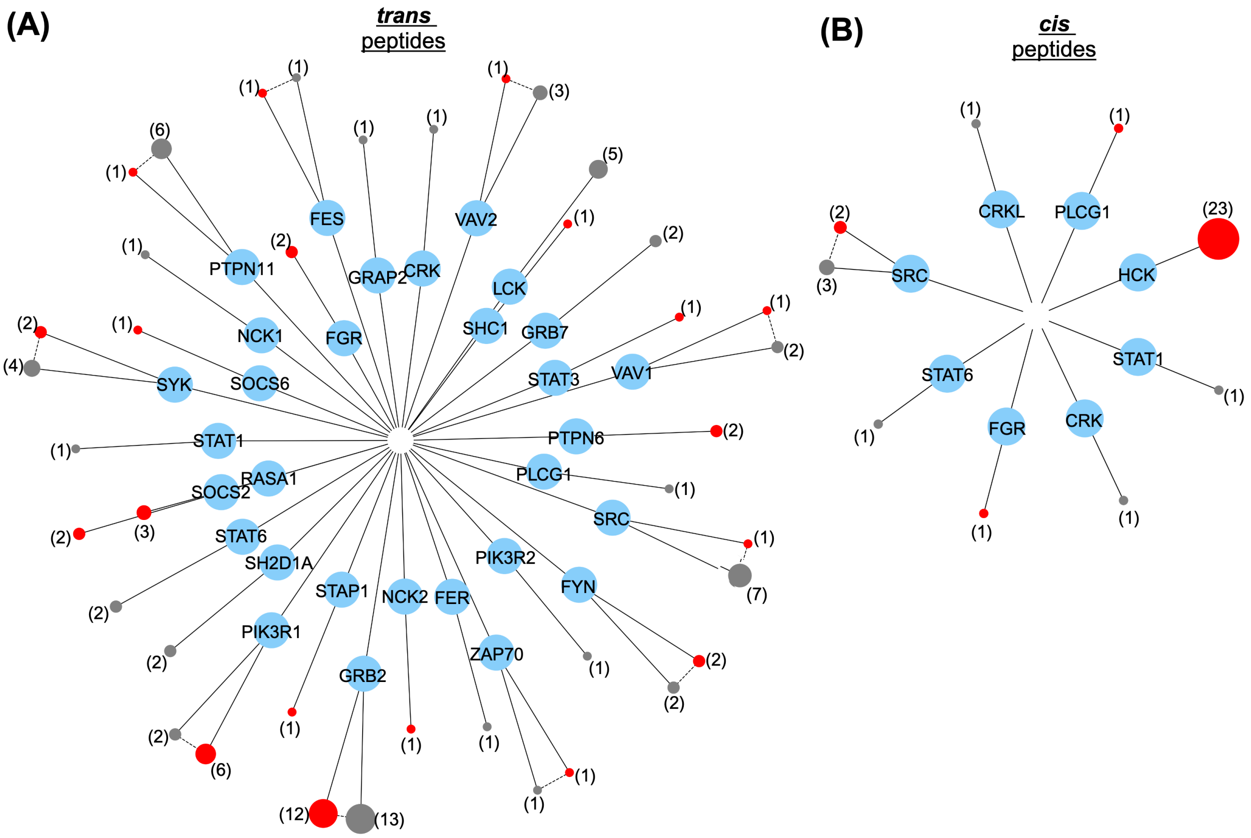


**Supplementary Figure 2. Structural coverage for PDB structures resolved for multi-entity protein complexes (SH2-pTyr).** This graph lays out the number of CoDIAC extracted PDB structures for the purpose of SH2-pTyr contact interface analysis. The number of structures capturing a SH2-pTyr interface for each gene are shown using this graph layout. The wild-type (grey nodes) and mutant (red nodes) structures for each gene are also shown. The sizes of the gene nodes indicate the number of structures and is denoted next to those nodes. **A**) The SH2 domain and pTyr peptide motif are on different entities that involve *trans*-like peptide interactions. (**B**) The number of *cis*-like peptide interactions of SH2-domains proteins is shown (the ligand and domain are on the same chain or there is a bivalent interaction).

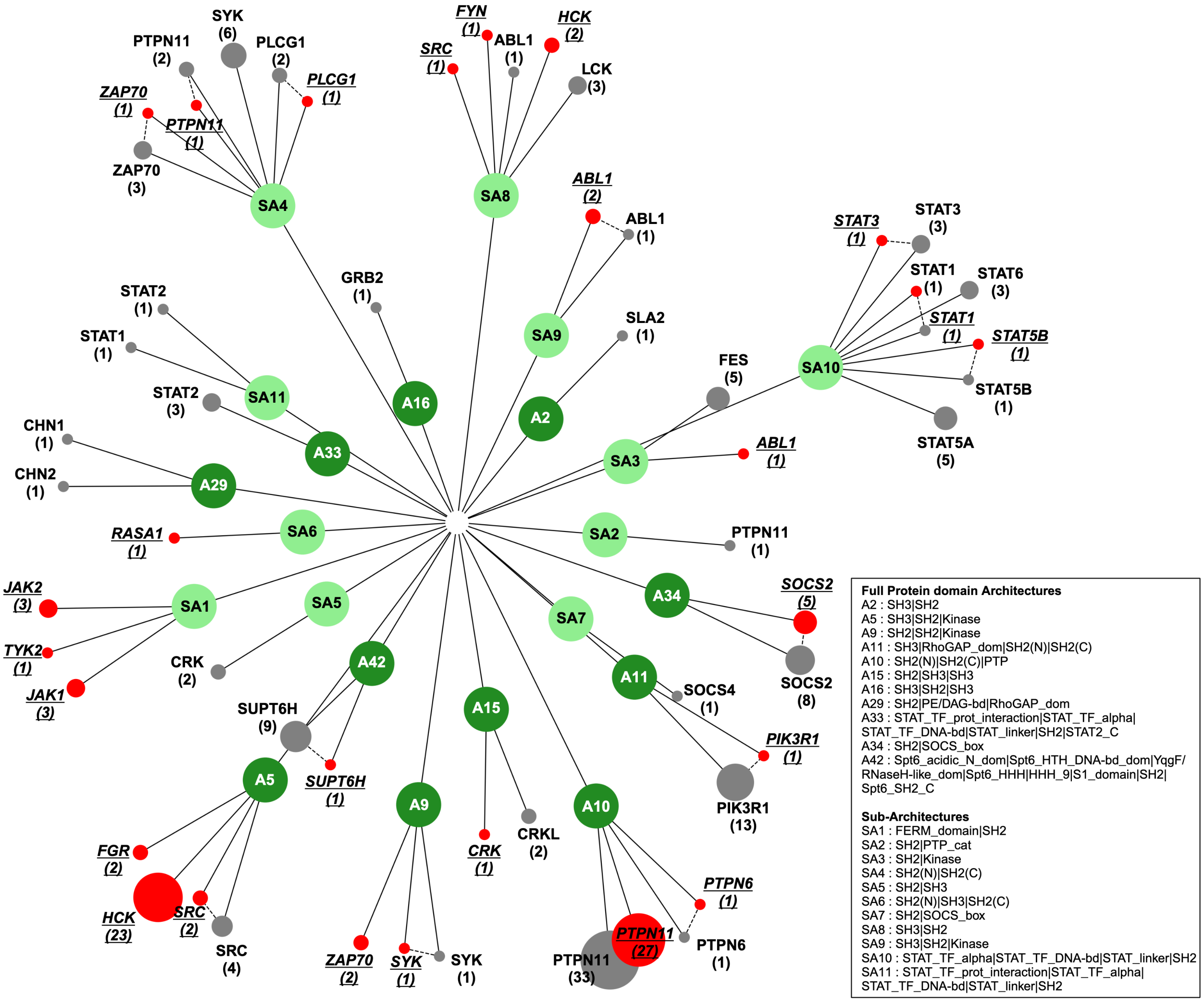


**Supplementary Figure 3. Structural coverage for PDB structures resolved for multi-domains.** This graph lays out the number of CoDIAC extracted PDB structures for the purpose of domain-domain contact analysis. The child nodes of this network represent wild-type (grey nodes) and mutant (red nodes) structures for each gene. The gene node sizes are indicative of the number of structures which is also denoted next to the gene nodes. Group architectures are from Fig. ??.

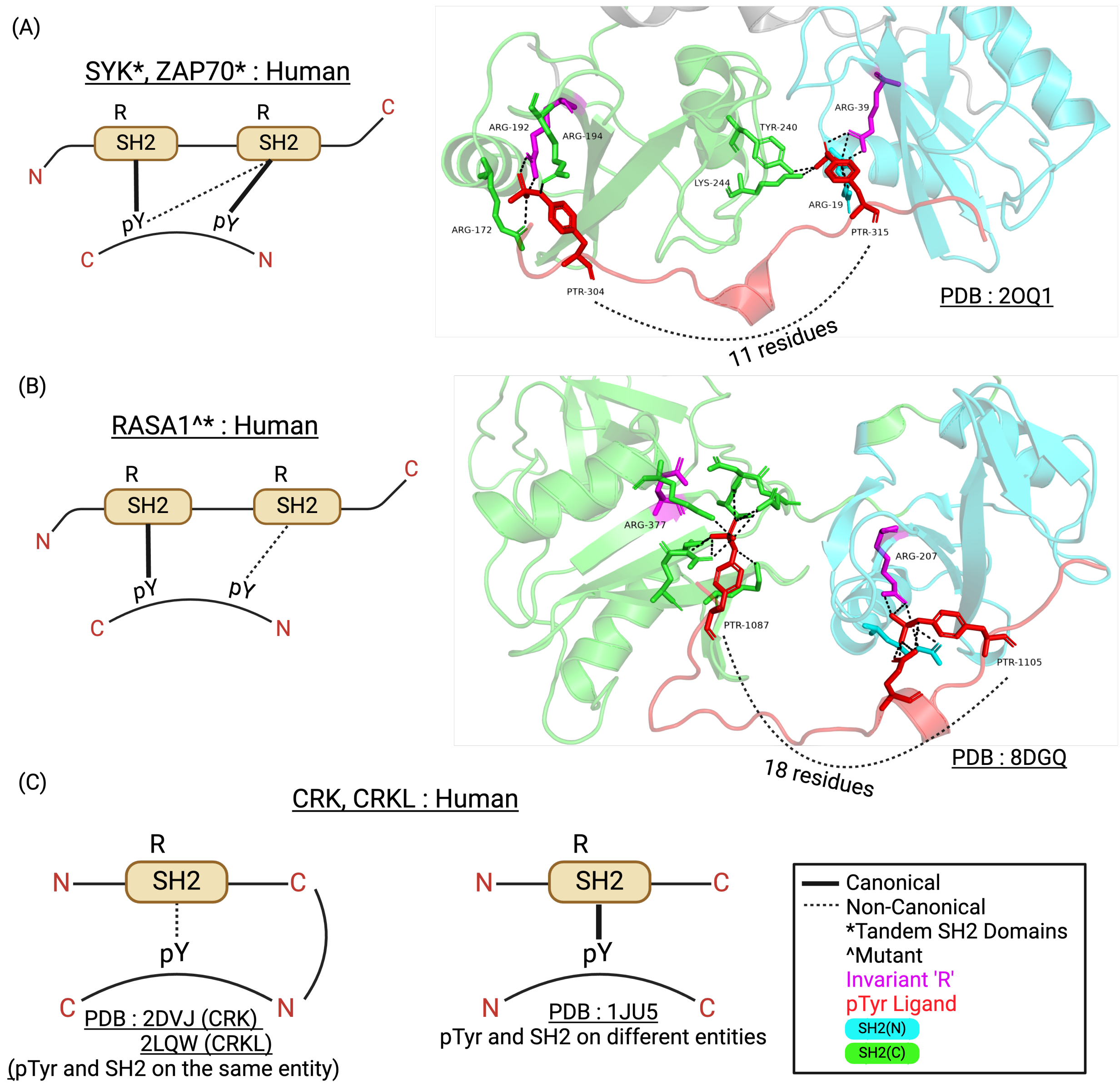


**Supplementary Figure 4. Examples of SH2 domain-ligand interactions that do not include the invariant arginine-pTyr interaction (i.e. non-canonical interactions).** (**A**) In SYK and ZAP70, one of the doubly phosphorylated tyrosines involve in only canonical binding and other pTyr interactions results in a canonical and non-canonical binding interface with each of the tandem SH2 domains. (**B**) In RASA1 that is a mutant structure, each of the tandem SH2 domains involve in either canonical or non-canonical binding with each of the double phosphotyrosine ligand. (**C**) In CRK and CRKL when the ligand is in cis-conformation with the SH2 domain, a non-canonical binding interface formation occurs and in a CRK structure forms a canonical interface when the ligand is tied in a trans conformation.

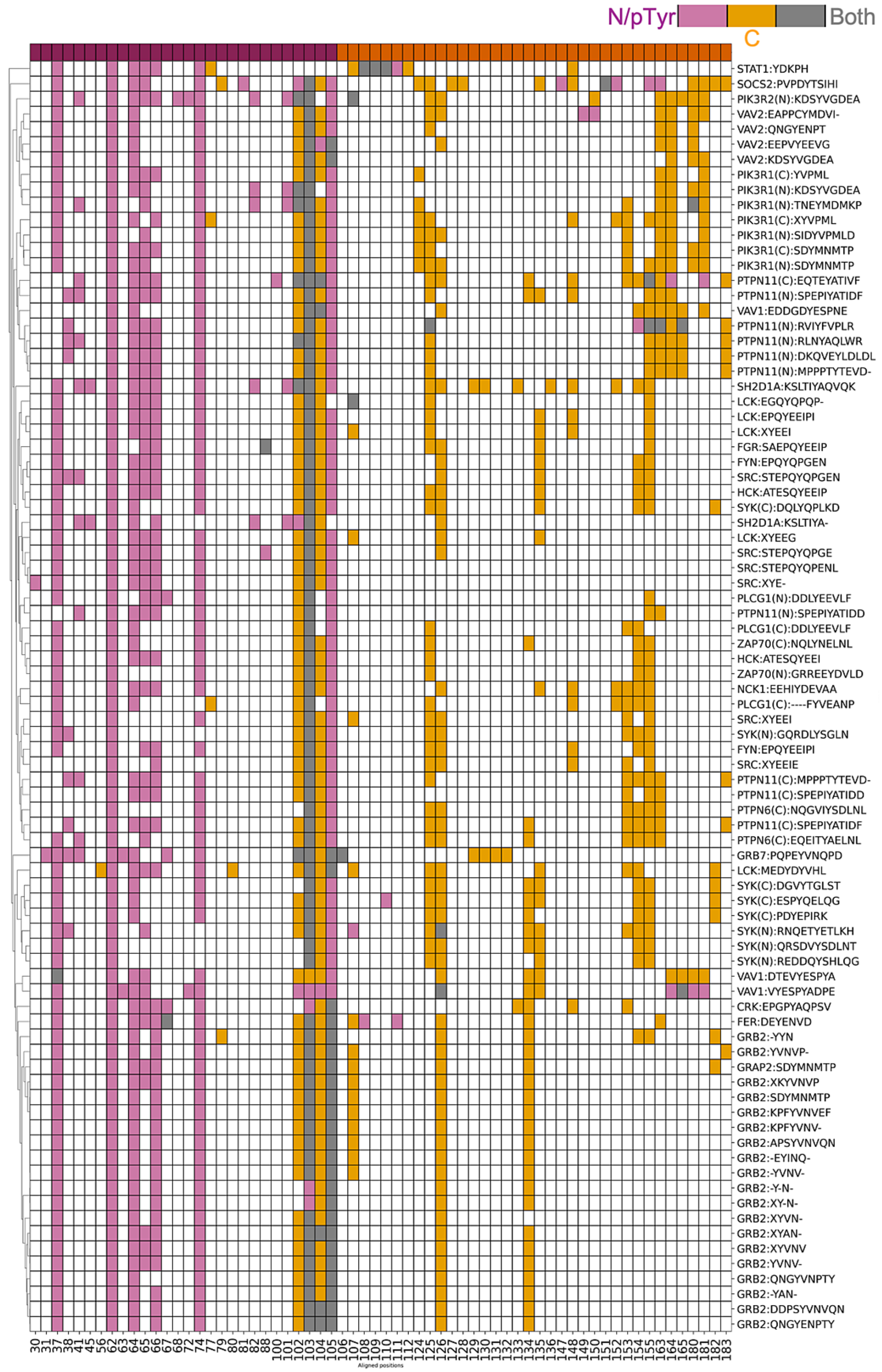


**Supplementary Figure 5. SH2-centric features for every unique SH2-pTyr peptide pair from PDB structures.** This clustermap is a representation of hierarchically clustered SH2-centric features (residues on SH2 domains that contact the ligand) that are extracted for 84 unique SH2-pTyr interfaces. Each contact position is colored encoded based of the section of the peptide (N/C/both) it has engaged with. The pink represent the N terminal (positions from N terminal up to pTyr), orange represents the C terminal (positions after pTyr residue) and grey for both sections.

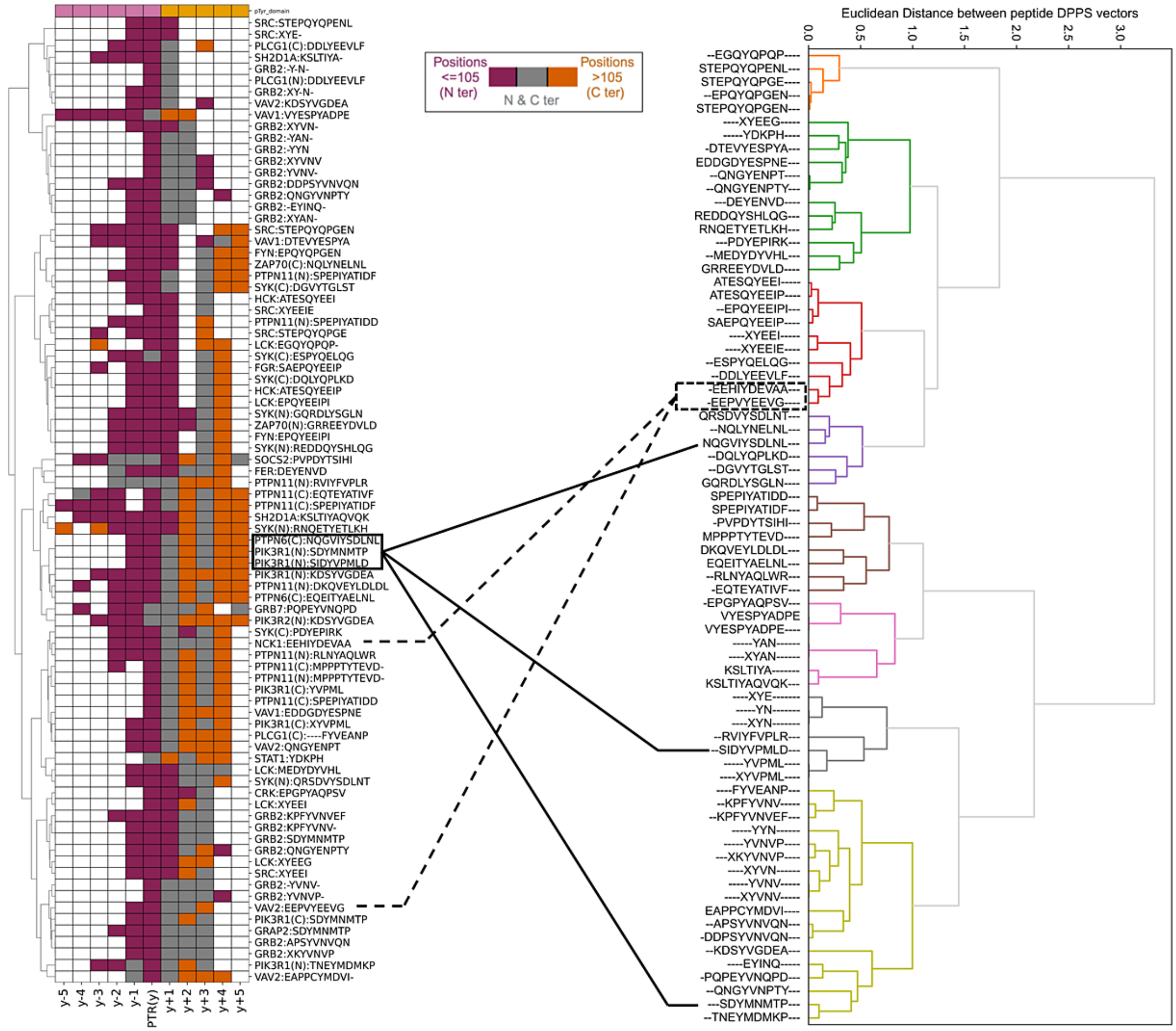


(Caption next page.)

**Supplementary Figure 5. Relationship between the contact maps and physiochemical property based grouping of pTyr peptides.** The ligand-centric contacts made by every unique pTyr peptide with an SH2 domain are clustered on the left and these contacts are color encoded based on the engagement with different sections of the SH2 domain. Euclidean distances calculated between pTyr peptides represented as DPPS (divided physicochemical property score) vectors are clustered on the right and these clustered groups capture peptides sharing similar physiochemical properties. Two cases are higlighted: 1) peptides sharing similar physiochemical properties make different set of contacts (dashed lines) and 2) ligands producing similar contacts do not necessarily correlate by their physiochemical characteristics.

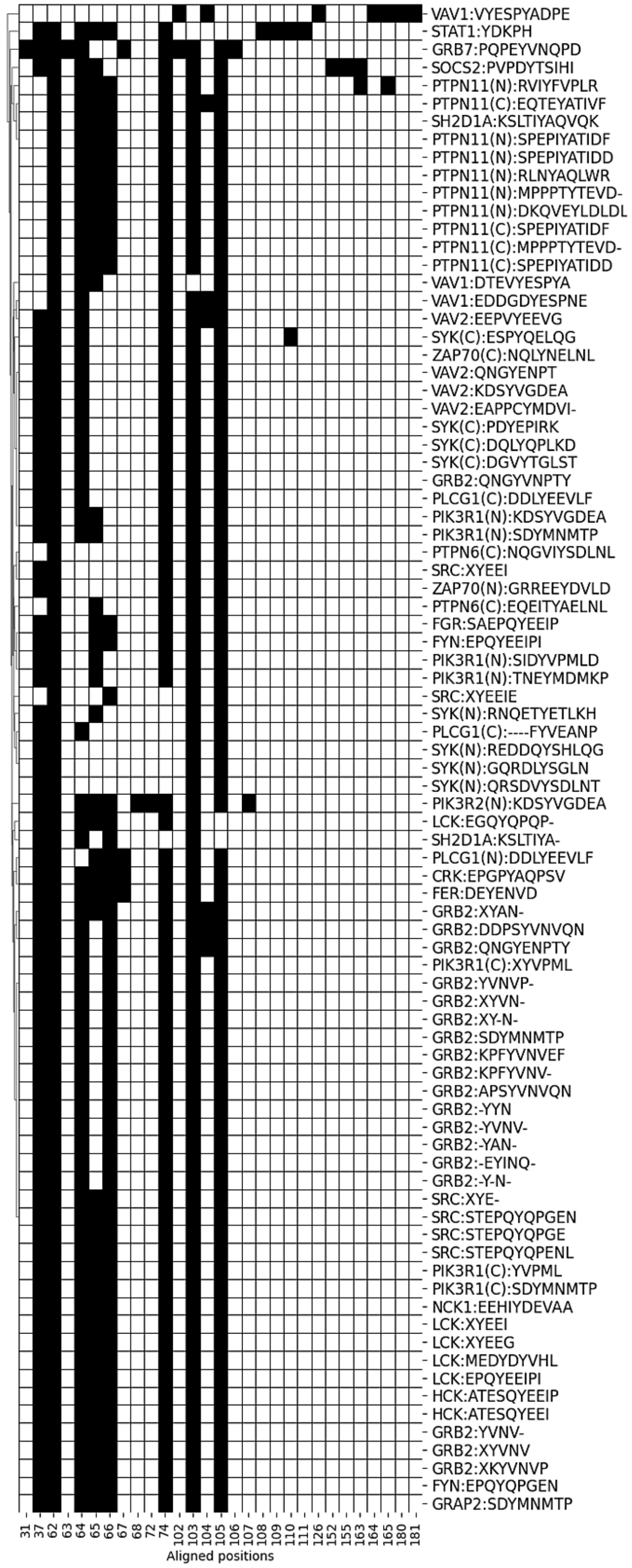


**Supplementary Figure 6. Specific contacts made by 'pTyr' with the residues on SH2 domains.**The binary matrix of SH2 alignment positions (x-axis) that contact the phosphotyrosine on the ligand shown on the y-axis. The binary matrix was then clustered (dendrogram on left). Position 105 represents the transition from the N-terminal to the C-terminal part of the SH2 domain.

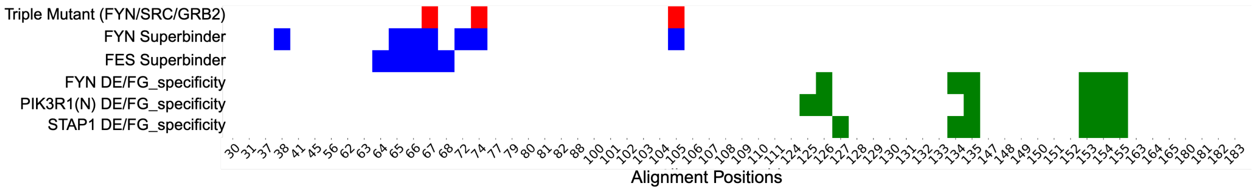


**Supplementary Figure 7. Projection of the superbinder mutations on the reference aligned SH2 sequence.** Three categories of superbinder mutations reported in Kaneko et al. [**?**] and Martyn et al. [**?**] were shown to affect either the affinity or specificity of the SH2 domains. We have placed each mutation set per protein on each line and indicated their effect (Superbinder indicates increase in binding affinity and specificity indicates it changed ligand specificity), relative to the alignment position of the SH2 domain family (x-axis). The triple mutant that has been used throughout several studies is on the top line.

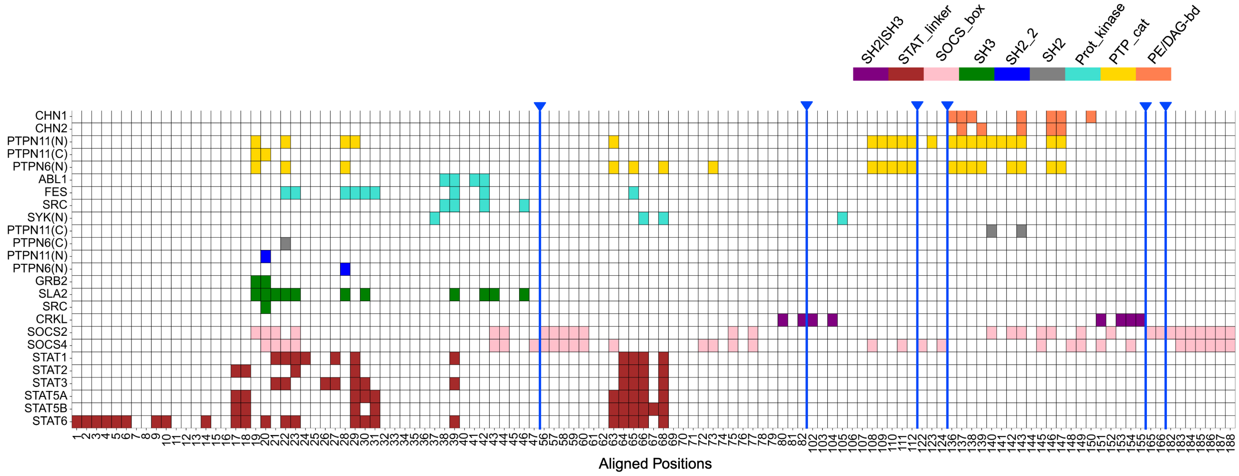


**Supplementary Figure 8. Heatmap of domain-domain contacts obtained from experimental structures.** The domain-domain contacts extracted using PDB structures are shown as a binary matrix (colored box indicates a contact for a protein between the domain of interest and the SH2 domain, on the alignment of the SH2 domain). Domain-centric contact maps for 21 SH2 domains and their engagement with nine other modular domains within the protein were extracted (color coded according to the domain interface). Alignment gaps between positions that are 5 amino acids or more are marked by blue arrows.

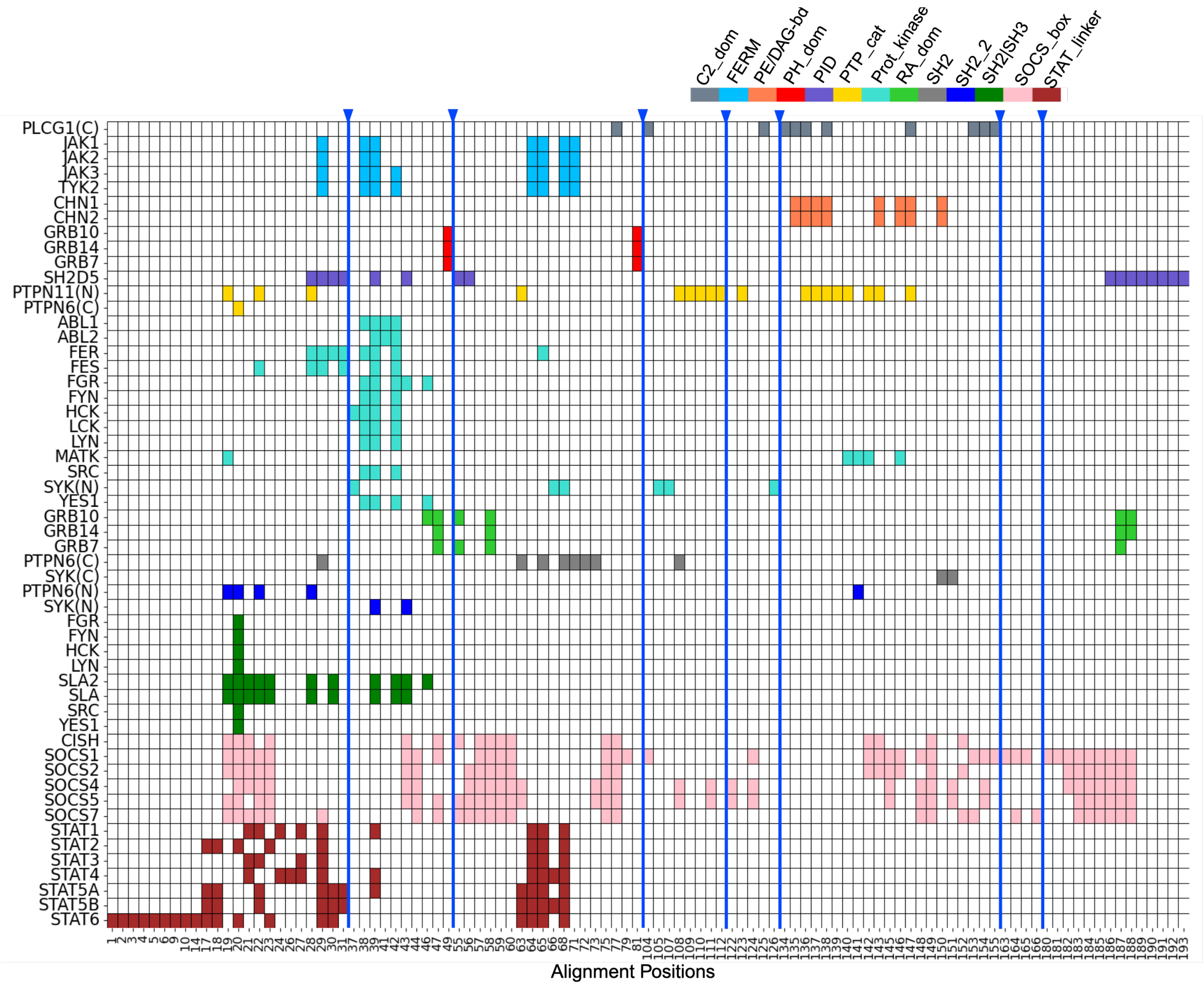


**Supplementary Figure 9. Heatmap of domain-domain contacts obtained from AlphaFold structures.** The domain-domain contacts extracted using 43 AlphaFold structures for 13 unique domain-domain interfaces are shown as a binary matrix (colored box indicates a contact for a protein between the domain of interest and the SH2 domain, on the alignment of the SH2 domain). The gaps on the aligned sequence of length 5 amino acids or more in this heatmap are indicated by blue arrows.

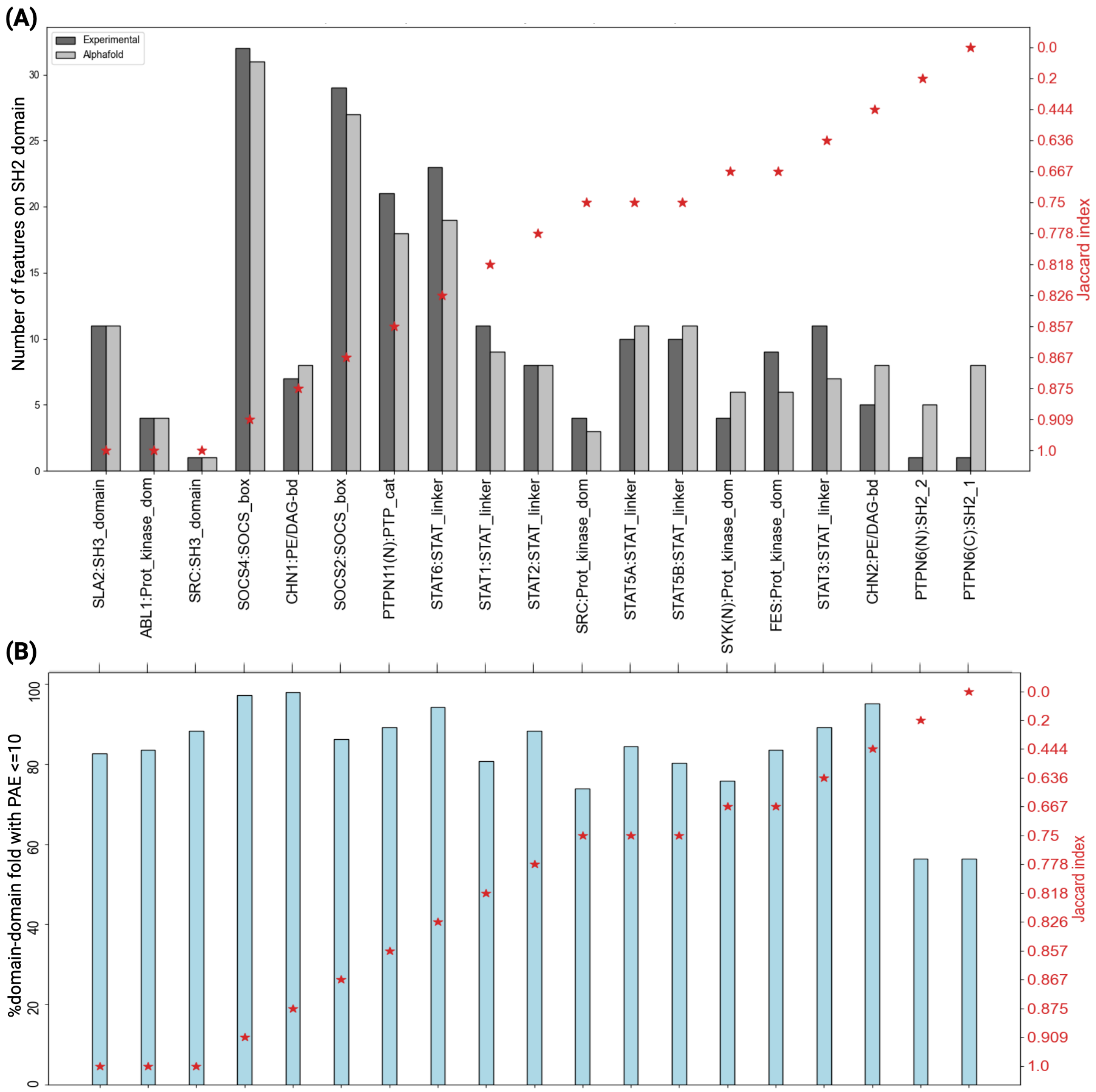


**Supplementary Figure 10. Comparison of domain-domain contacts extracted from experimental and predicted structures.** (**A**) The domain-domain interfaces for which we extract data from both experiments and predicted structures are compared and measured for similarity of the contacts using Jaccard index (red stars, right-axis). Bars for each domain-domain interface depict the total number of contacts from experimental and predicted structures (left axis). (**B**) AlphaFold PAE score (left axis) for each comparison found in panel A is shown, along with the corresponding JI value (red). Correlation between experimental and predicted structures was used to identify meaningful PAE scores for inclusion.

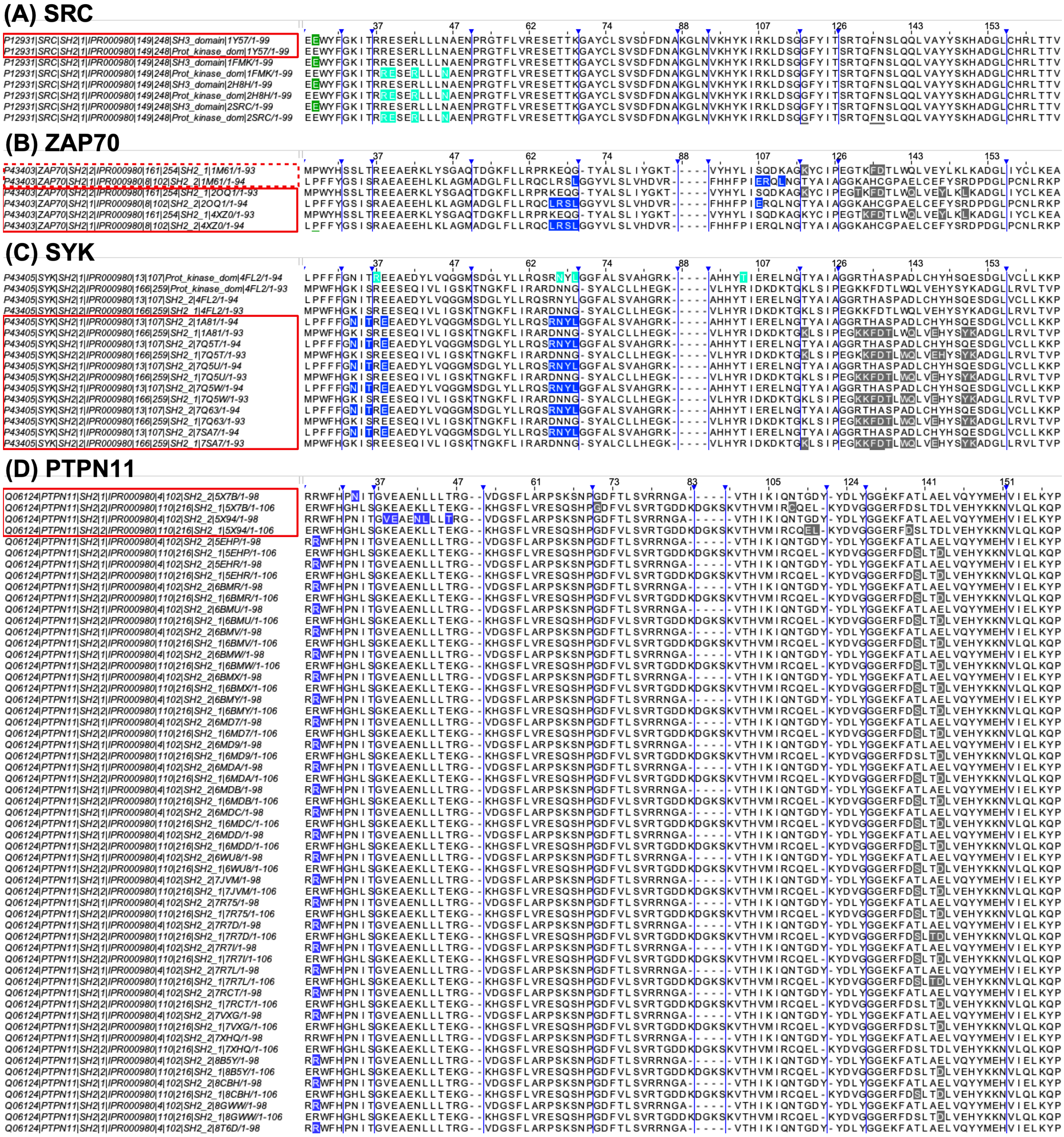

**Supplementary Figure 11. Analysis of domain-domain interfaces for partial versus complete experimental structures.** Alignments are shown for 4 SH2 domains: (**A**) SRC (**B**) ZAP70 (**C**) SYK (**D**) PTPN11, where extracted interfaces (highlighted in a color associated with the domain interfaces found in fig. **??** varied significantly based on the structure. Red boxes are placed around structures with low RMSD.

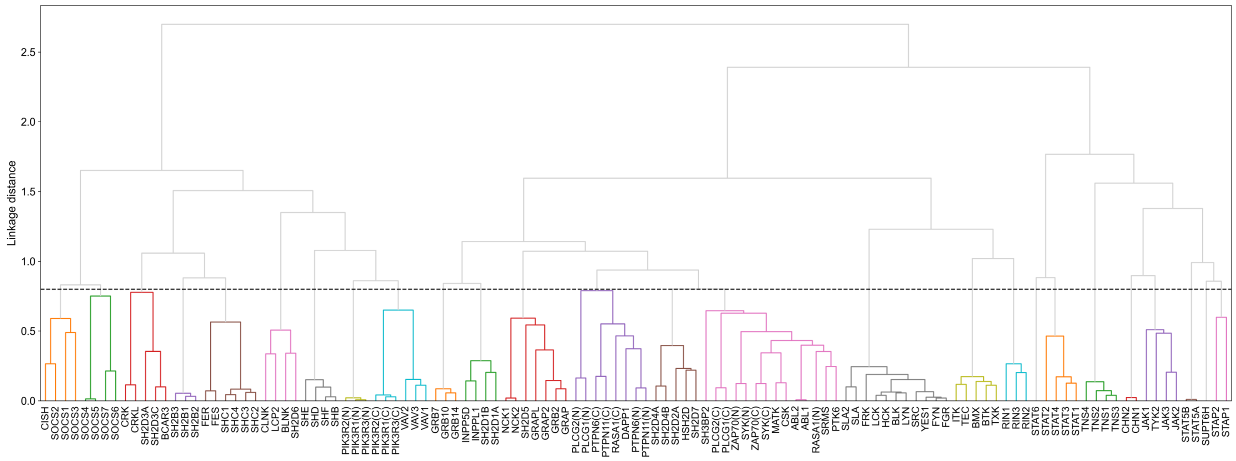


**Supplementary Figure 12. Dendrogram to illustrate the grouping the 119 human SH2-domains using relative evolutionary distances.** The phylogenetic tree is shown, using a distance-based Neighbor-joining method. The 'SH2ome' is divided to 26 clusters at an optimum threshold of 0.8 linkage distance and every colored set of leaves of the tree represent each of these 26 clusters.

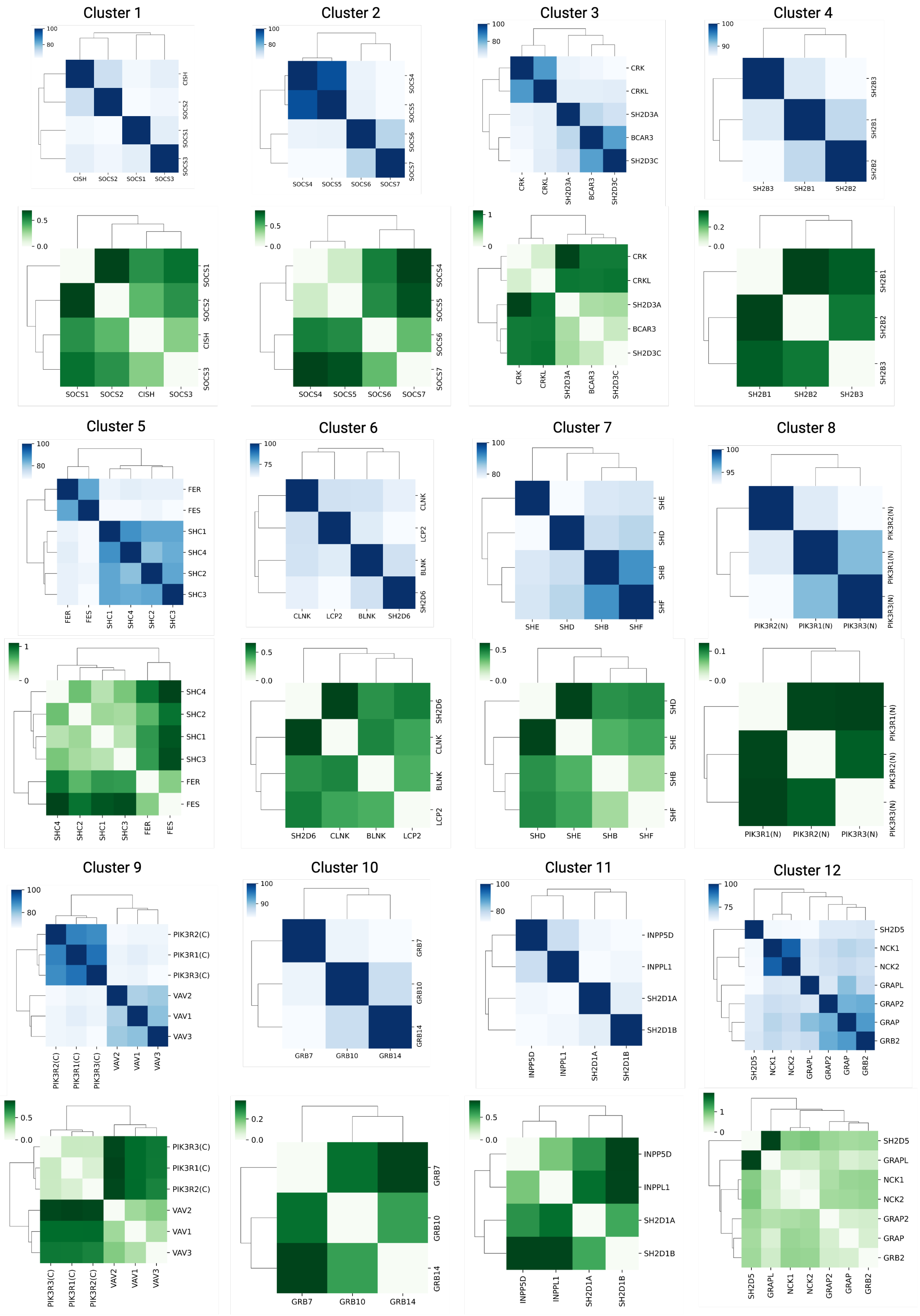

Cluster 1
100
80
CISH
SOCS2
SOCS1
SOCS3
0.5
0.0
Cluster 2
100
80
SOCS4
SOCS5
SOCS6
SOCS7
0.5
0.0
Cluster 3
100
80
CRK
CRKL
SH2D3A
BCAR3
SH2D3C
1
0
Cluster 4
100
90
SH2B3
SH2B1
SH2B2
0.2
0.0
Cluster 5
100
80
FER
FES
SHC1
SHC4
SHC2
SHC3
1
0
Cluster 6
100
75
CLNK
LCP2
BLNK
SH2D6
0.5
0.0
Cluster 7
100
80
SHE
SHD
SHB
SHF
0.5
0.0
Cluster 8
100
95
PIK3R2(N)
PIK3R1(N)
PIK3R3(N)
0.1
0.0
Cluster 9
100
80
PIK3R2(C)
PIK3R1(C)
PIK3R3(C)
VAV2
VAV1
VAV3
0.5
0.0
Cluster 10
100
90
GRB7
GRB10
GRB14
0.2
0.0
Cluster 11
100
80
INPP5D
INPPL1
SH2D1A
SH2D1B
0.5
0.0
Cluster 12
100
75
SH2D5
NCK1
NCK2
GRAPL
GRAP2
GRAP
GRB2
1
0

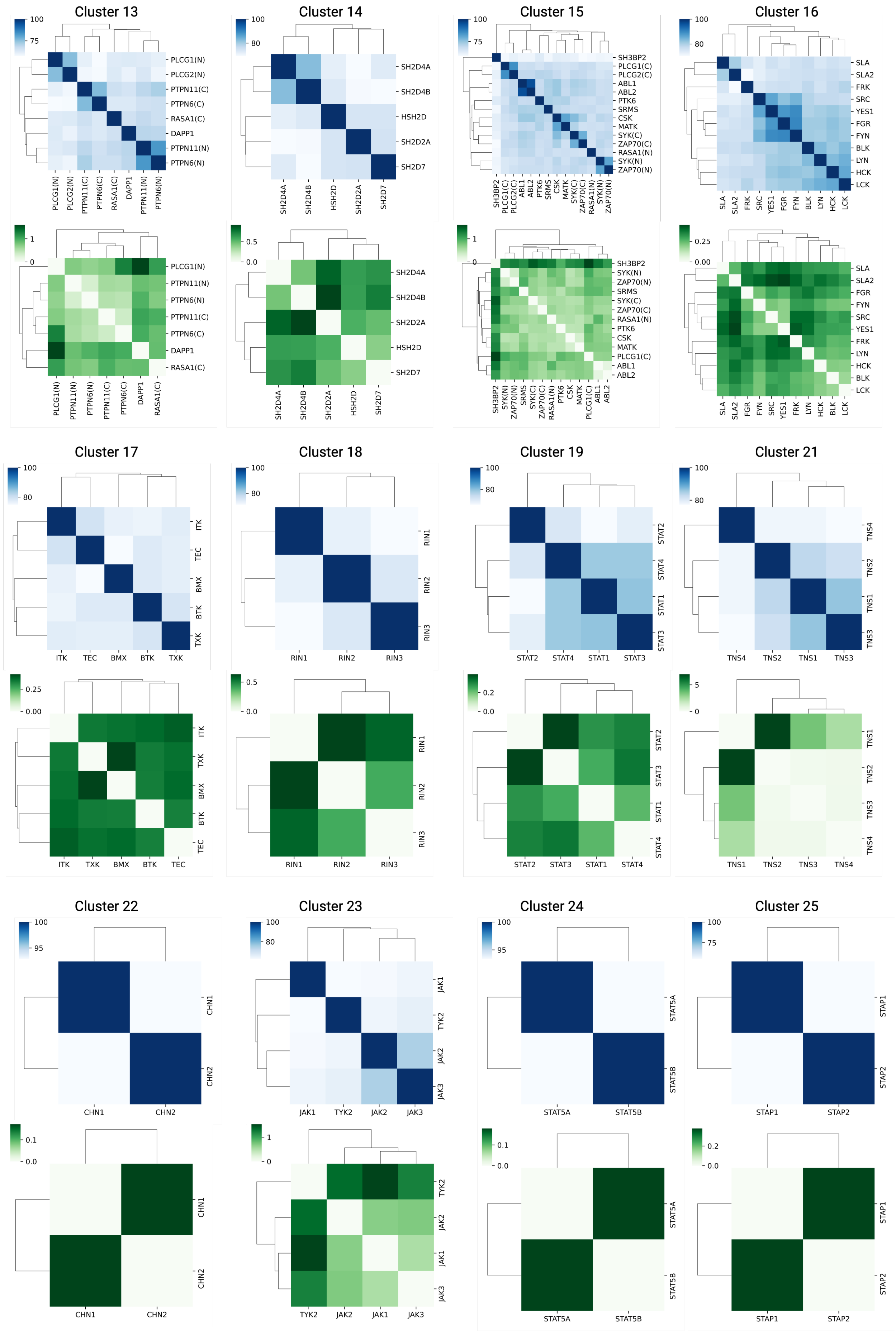


**Supplementary Figure 13. Sequence and structure similarity for each of the classified SH2 domain clusters.**

**Sequence and structure similarity for each of the classified SH2 domain clusters (clusters appearing on prior two pages.** Each of the 26 clusters is further sub-divided to find intra-clustered homologous proteins. The sub-clustering was performed using sequence identity scores (blue heatmaps) and RMSD values (green heatmaps) for sequence and structure comparisons respectively.

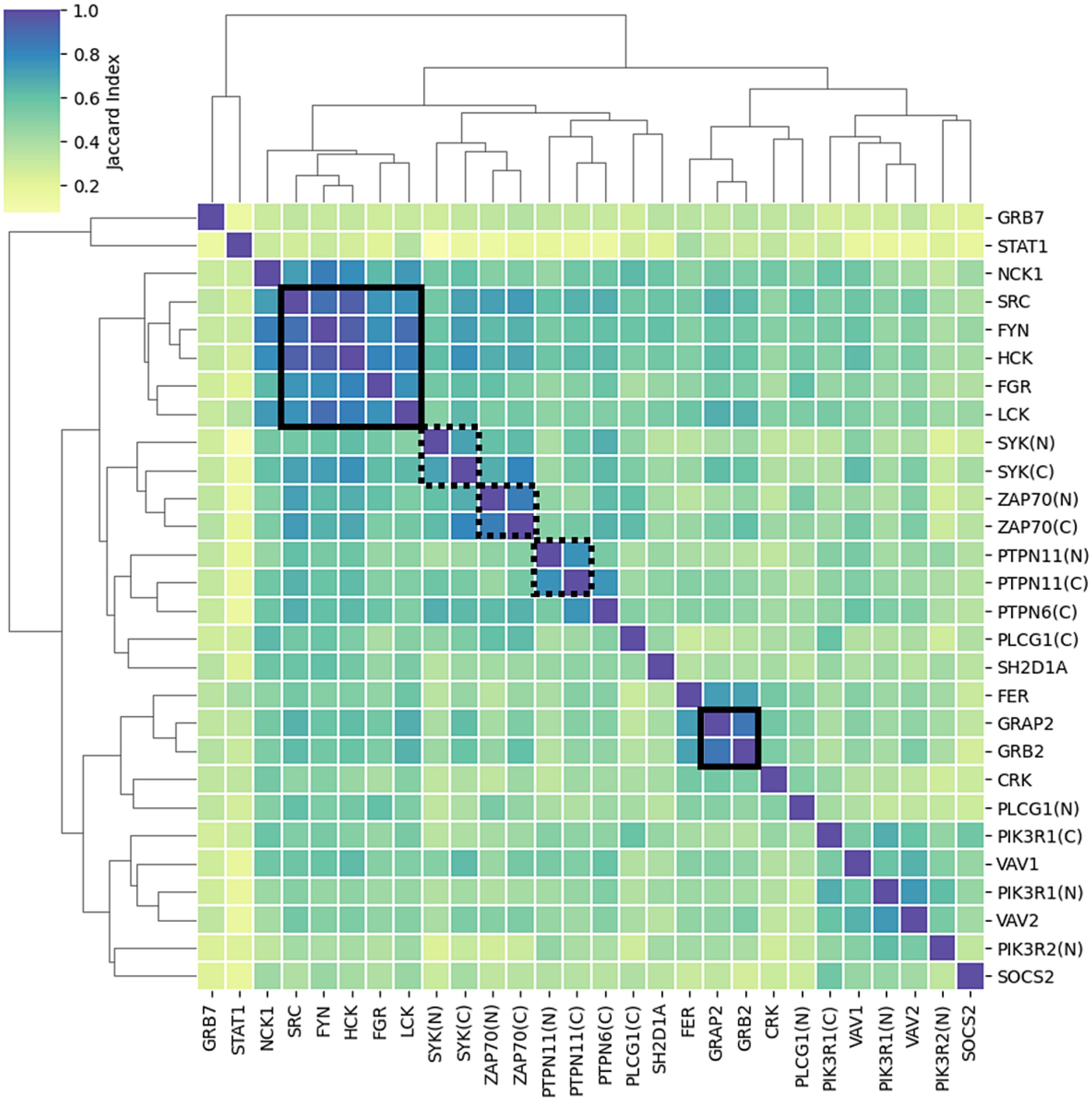


**Supplementary Figure 14. Comparison of experimentally determined ligand contact maps across clustered proteins.** The heatmap, based on the Jaccard index comparison of ligand contacts extracted between all pairs of SH2 domains available, is hierarchically clustered. High similarity groups are outlined (solid boxes around single SH2 domain containing proteins and dashed around tandem domains from the same protein).

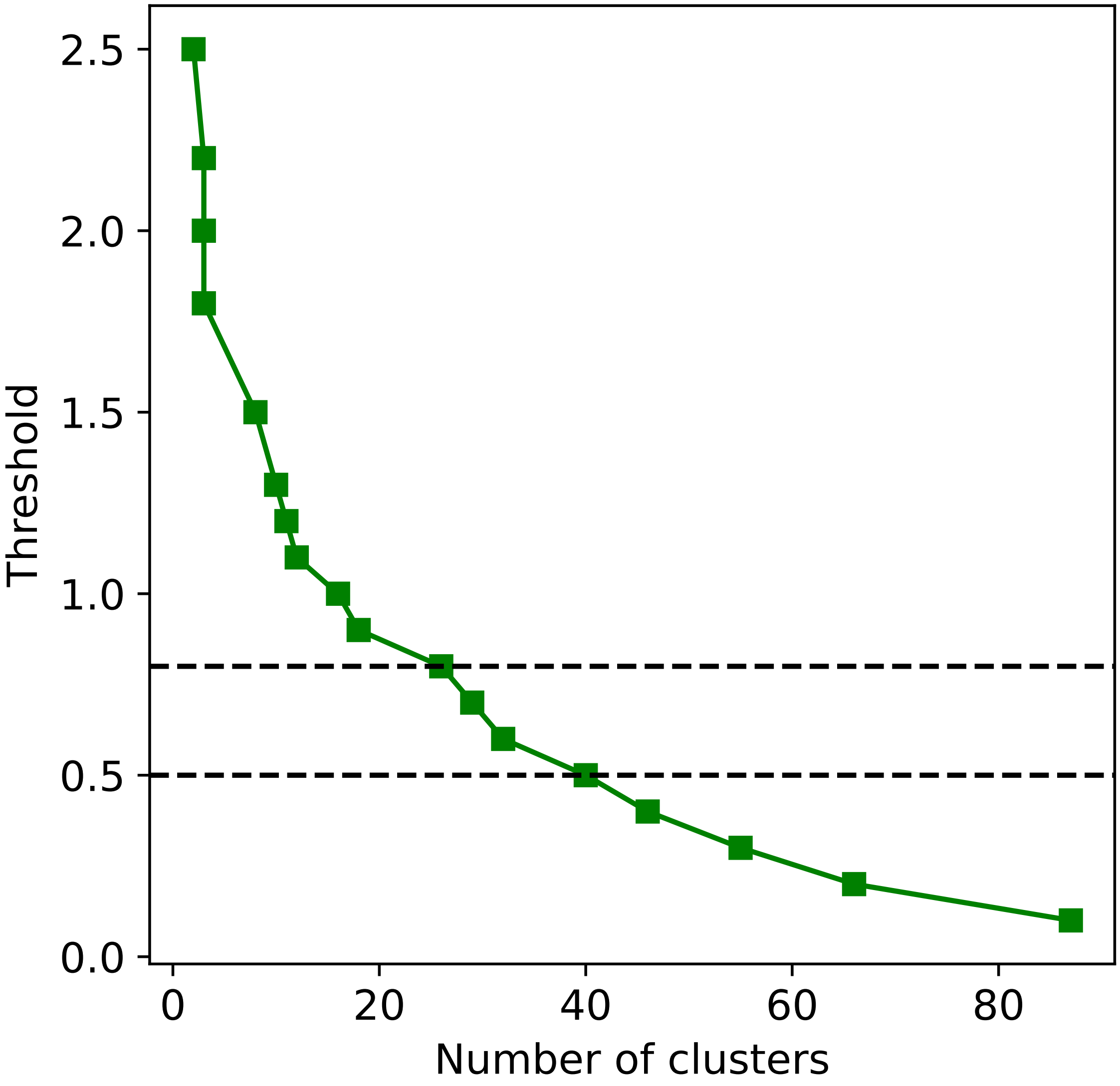


**Supplementary Figure 15. Optimum threshold for SH2 domain protein classification using elbow method.** The phylogenetic tree cutoff (y-axis) versus the number of clusters produced (x-axis) of fig. **??** is shown here for identification of possible clusters. We have given the cluster membership of a 0.8 (26 clusters) and 0.5 (40 clusters) cutoff in Data File S1A and S1D, respectively.

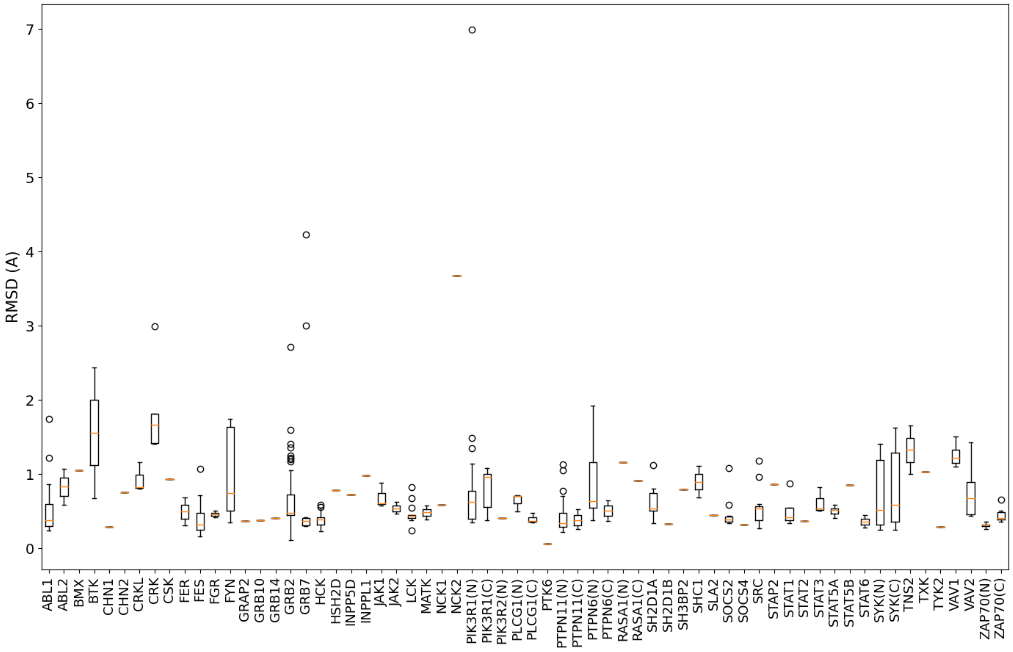


**Supplementary Figure 16. Structural comparison of experimentally and predicted SH2 domain structures.**The RMSD (angstroms, y-axis) was computed for every experimental structure available to the predicted AlphaFold structure. SH2 domains are sorted alphabetically.

**Supplementary Tables**

**Supplementary Table 1. Statistical testing of acidic amino acids in the -1 ([DE]y) and +1 (y[DE]) positions.** FDR corrected p-values are reported for each Fisher's exact test, along with the number of peptides that matched the motif in the background (n) and foreground from Martyn et al. [**?**]. Structure extracted indicates if there is a complementary structure data and FDR values less than 0.05 are in bold.

| Structure extracted | | n | [DE]y | [DE]y FDR | y[DE] | y[DE] FDR |
|---|---|---|---|---|---|---|
| | background | 5192 | 1095 | | 1334 | |
| (neg control) | Ti-Zr-IMAC | 1801 | 398 | 5.56E-01 | 443 | 0.524333 |
| Yes | sNck1-S | 1021 | 209 | 7.42E-01 | 487 | **4.93E-21** |
| Yes | sLck-S | 589 | 107 | 3.02E-01 | 268 | **1.50E-11** |
| Yes | Src-wt | 511 | 120 | 4.67E-01 | 229 | **8.87E-10** |
| Yes | sVAV3-S | 868 | 152 | 8.99E-02 | 338 | **2.25E-08** |
| Yes | sPTN11_N | 100 | 17 | 5.56E-01 | 6 | **7.71E-05** |
| Yes | sPTN6_C | 383 | 51 | **6.14E-03** | 57 | **1.59E-04** |
| Yes | PIK3R1_N | 98 | 49 | **6.59E-05** | 39 | **3.60E-02** |
| Yes | PIK3R2_N | 100 | 45 | **4.07E-04** | 39 | **4.37E-02** |
| Yes | CRKL-wt | 151 | 15 | **7.58E-03** | 47 | 3.34E-01 |
| Yes | sGrb2-S | 577 | 72 | **1.37E-04** | 146 | 9.23E-01 |
| No | sFes-S | 625 | 141 | 5.56E-01 | 438 | **9.28E-44** |
| No | sAbl1-S | 1055 | 172 | **7.58E-03** | 411 | **1.41E-09** |

**Caption for Data S1. SH2 domain clusters and analysis.** Clusters (Cluster ID) and Subclusters (Sub ID) were produced by evolutionary distance clustering with different cutoffs and assessed for their ability to group SH2 domains based on ligand binding and domain-domain interactions. (**A**) Evolutionary distance-based clustering produces 26 clusters for a linkage threshold 0.8. (**B**) Non-tandem SH2-domain protein classification. The 26 clusters obtained using the evolutionary distances is the basis for 'SH2ome' classification. One of the clusters comprises of only tandem SH2-domains (highlighted in red) and this sub-clustered separately along with other tandem SH2-domain proteins. The remaining 25 cluster groups were further grouped into smaller clusters to find the closest neighbors. The comparisons for intra-clustered homologues using primary amino acid sequence (sequence identity) and structure (RMSD) to validate the evolutionary-based outcomes is indicated in this table. If the intra-clustered homologues using sequence identity and structure similarity scores were identical to the evolutionary based sub-grouping, we indicated those by '1'. The clusters comprising of less than two proteins, then we indicate those by '-'. Similarity between the contacts (Jaccard index greater than 0.5) formed across proteins within sub-clustered is also indicated as '1' and this is used to validate our grouping outcomes. We find that the contact similarity in clusters containing tandem SH2 domain proteins do not agree with the grouping based on evolution/sequence/structure and hence grouped them differently from the rest of the non-tandem SH2-domain proteins and these are indicated as '0'. If the sub-clustering outputs agree in all the three categories, then we keep those sub-clusters as is (differentiated by dashed lines) otherwise we group them into one cluster (for example cluster 1). The cluster and sub-cluster numbering are designated in the last two columns. For contact projection, we first use the sub-clusters to propagate contacts on to SH2 domains and on unavailing any extracted protein interfaces, we extend our search to the cluster. The final 42 sub-clusters listed here were employed for contact projection across domain-domain and ligand binding interfaces. (**C**) Tandem SH2-domain proteins classification used for contact projection across ligand and domain-domain binding interfaces. For ligand contact interface projection, the (N) and (C) SH2 domains of the same protein are grouped together since their feature similarity is significantly higher than how the evolution and sequence would direct (for example N terminals of PTPN11 and PTPN6 together). For domain-domain interface contact projection, we follow the evolutionary based grouping. We employed these cluster grouping for contact map projection within tandem SH2 domain proteins. (**D**) Evolutionary distance-based clustering produces 40 clusters for a linkage threshold 0.5.

**Caption for Data S2. PAE of AlphaFold predictions and RMSD analysis of PDB structures.** (**A**) PAE is reported for the % of the predicted structure pairwise domains being considered that are less than 10 angstroms in error. These guided decisions for exclusion of a predicted structure for domain-domain interfaces. (**B**) Structural comparison of SH2 domains using experimentally determined and predicted structures for all domains with more than one structure and with AlphaFold predicted structures (that meet PAE range). PDB or AlphaFold predictions deemed erroneous, compared to remaining, are eliminated and highlighted in red.

**Caption for Data S3. Post-translational modification reports.** Sheets are indicated for the PTM analysis (phosphorylation of serine/threonine, phosphorylation of tyrosine, N6-acetyllysine, and ubiquitination). The data are sorted by the common alignment position and the number of conserved modifications found in that position. Groups of note have been colored and text boxes call out specific hypotheses. Data table headers are: header (the fasta header referring to the specific SH2 domain); Uniprot_ID (Uniprot ID of the protein); SH2 number (for tandem domains, this is 1 for N-term and 2 for the C-terminal domain); protein_residue (residue number in the full length protein of Uniprot ID); feature_position (numbering relative to start of the SH2 domain); alignment_postion (position in the reference alignment used throughout this work); number_mods_at_position (total number of this type of modification at that alignment position across all SH2 domains); number_conserved_ligand_residues_nearby (in the window of +/-2 amino acids, how many ligand biding positions are nearby); number_residues_from_conserved_ligand (how far away is the closest conserved ligand binding position, 0 for it is a ligand position); domain_interaction (the other domain, if applicable, that interacts with this position); Literature citations (citations related to that PTM).

**Caption for Data S4. Mutation report.** (**A**) Evaluation of mutations based on where they fall at binding interfaces and on or near conserved PTMs. Similar to data S3, data are sorted based on most number of mutations found at that alignment position and groups of interest are colored. Data table headers are: header (the fasta header referring to the specific SH2 domain); Uniprot_ID (Uniprot ID of the protein); gene_name (name of gene); SH2 number (for tandem domains, this is 1 for N-term and 2 for the C-terminal domain); protein_residue (residue number in the full length protein of Uniprot ID); feature_position (numbering relative to start of the SH2 domain); alignment_position (position in the reference alignment used throughout this work); number_mutations_at_position (total number of mutations at that alignment position across all SH2 domains); number_conserved_ligand_residues_nearby (in the window of +/-2 amino acids, how many ligand biding positions are nearby); number_residues_from_conserved_ligand (how far away is the closest conserved ligand binding position, 0 for it is a ligand position); domain_interaction (the other domain, if applicable, that interacts with this position); PTMs (indicates if this mutation directly occurs on a residue known to be modified, detailing the modification). (**B**) Mutations are classified according to the effect of the substitution by charge or polarity change for mutations found at ligand or domain-domain contact interfaces.. The Gene, the mutation (full length protein residue) are given, along with the classification.
